# Interaction with C21ORF2 controls the cellular functioning of the NEK1 kinase

**DOI:** 10.1101/2022.08.31.505651

**Authors:** Mateusz Gregorczyk, Graziana Pastore, Pawel Lis, Sven Lange, Frederic Lamoliatte, Thomas Macartney, Rachel Toth, Fiona Brown, James Hastie, Daniel Durocher, John Rouse

**Affiliations:** MRC Protein Phosphorylation and Ubiquitylation Unit, School of Life Sciences, Wellcome Trust Biocentre, University of Dundee, DD1 5EH, UK; The Lunenfeld-Tannenbaum Research Institute, Mount Sinai Hospital Rm 1073, 600 University Avenue, Toronto, Ontario, M5G 1X5 Canada; Department of Molecular Genetics, University of Toronto, Toronto, Ontario M5S 1A8 Canada.

**Keywords:** NEK1, C21ORF2, kinase, ALS, SMD, ciliogenesis, DNA repair

## Abstract

NEK1 is a pleiotropic protein kinase implicated in mitosis, ciliogenesis and DNA repair but little is known about its regulation or targets. Its relevance for human health is underscored by the association of *NEK1* mutations with human diseases including axial spondylometaphyseal dyplasia (SMD) and amyotrophic lateral sclerosis (ALS). Mutations in the *C21ORF2* gene are associated with a similar pattern of human diseases, suggesting close functional links with *NEK1*. Here we report that in unperturbed, untransformed cells, endogenous NEK1 and C21ORF2 form a tight complex that does not appear to contain other proteins. A small acidic domain “CID: C21ORF2 interaction domain” at the C-terminus of NEK1 is necessary and sufficient to interact with C21ORF2, and pathogenic mutations in this region disrupt the complex. AlphaFold modelling predicts with high confidence an extended binding interface between a leucine-rich repeat (LRR) domain in the N-terminal half of C21ORF2 and a stretch of the NEK1-CID; mutating residues mediating electrostatic interactions within this interface disrupts the NEK1-C21ORF2 interaction. This model also explains why pathogenic mutations disrupt the complex. We go on to show that the kinase activity of NEK1 and its interaction with C21ORF2 is critical for NEK1 function in cells. These data reveal C21ORF2 as a regulatory subunit of NEK1, illuminating our understanding of how this kinase is regulated and NEK1-C21ORF2-associated diseases.

## Introduction

It is over three decades since the NEK1 protein kinase was discovered, and although it is essential for preventing a range of debilitating diseases, remarkably little is known concerning the molecular mechanisms underlying the roles and regulation of this enzyme. NEK1 is one of eleven members of the NEK family of protein serine/threonine kinases related to the NIMA (“<u>n</u>ever in <u>m</u>itosis”) kinase from *Aspergillus nidulans* (Letwin et al, 1992; O’Regan et al, 2007; Schultz & Nigg, 1993). NIMA is important for controlling mitotic entry (Morris, 1975; Oakley & Morris, 1983), and cells from NEK1-deficient mice show disordered mitoses, errors in mitotic chromosome segregation and cytokinesis (Chen *et al*, 2011; O’Regan *et al*., 2007). Moreover, abnormal organization of the meiosis I spindle and faulty chromosome congression have been observed in gametes from these mice (Brieno-Enriquez *et al*, 2017). Although NEK1 is clearly involved in mitotic function the underlying mechanisms and the relevance of kinase activity remain to be determined.

NEK1 localises to centrosomes which, in addition to organizing the mitotic spindle, nucleate the formation of primary cilia at the cell surface (Mahjoub *et al*, 2005). Primary cilia are single, thin projections found on the surface of most mammalian cells, predominantly on non-dividing or quiescent cells although they can also occur in G_1_ phase of proliferating cells (Satir & Christensen, 2007; Zimmerman, 1898). Cilia are tethered at their base by the mother centriole of the centrosome, from which an array of microtubules protrude, sheathed in plasma membrane, to provide scaffolding for the axonemal projection. These projections are sealed off at the base from the cytosol, with cargo trafficked in and out by dedicated intra-flagellar transport proteins. In this light, cilia are highly enriched for subsets of signalling proteins, acting as major hubs for Hedgehog (Shh) signalling critical for multiple aspects of cell function (Wheway *et al*, 2018). Mouse and human cells lacking functional NEK1 show pronounced defects in ciliogenesis and cilium function (Evangelista *et al*, 2008; Shalom *et al*, 2008). Furthermore, NEK1-deficient mice (*kat* and *kat2j*) show display symptoms typically associated with ciliopathies, diseases caused by ciliary defects, including polycystic kidney disease (PKD) (Chen *et al*, 2014; Mahjoub *et al*., 2005; Upadhya *et al*, 2000; Vogler *et al*, 1999). Even though NEK1 seems to play important roles in ciliogenesis, the underlying mechanisms remain to be determined and it is not yet clear if kinase activity is important.

NEK1–deficient mice show a range of defects beside kidney disease, including facial dysmorphism, dwarfism, cystic choroid plexus, male sterility, and anaemia (Janaswami *et al*, 1997; Upadhya *et al*., 2000). Sterility and anemia are often associated with defects in DNA repair, and in this light NEK1 has been implicated in homologous recombination (HR), a highly conserved pathway important for repairing double-strand breaks (DSB) in proliferating cells (Spies *et al*, 2016). HR uses the intact chromatid present in cells in late S- and G_2_-phases to direct repair of DNA breaks. DSB arising in S- and G_2_-phases are resected through the action of nucleases and helicases resulting in formation of 3’ single-stranded (ss) overhangs (Bonilla *et al*, 2020; Wright *et al*, 2018). RAD51 coats the ssDNA to form a nucleoprotein filament which searches for sequence homology in the intact chromatid, invades the homologous duplex and directs annealing to the homologous sequences. The invading strand is then extended using a form of non-canonical DNA synthesis (Bonilla *et al*., 2020; Saredi & Rouse, 2019). The final stages of HR can result in the formation of structures that entangle the two chromatids, which are enzymatically removed to complete repair (Lilley, 2017). Depletion of NEK1 from HeLa cells was reported to cause near-complete abrogation of DSB-induced HR; this appeared to stem from loss of NEK1-mediated phosphorylation of the RAD54 ATPase on Ser 572, which was in turn required for RAD54-mediated unloading of RAD51 from DSB repair sites (Spies *et al*., 2016). In principle, failure to unload RAD51 could prevent completion of HR by preventing extension of the invading strand. However, RAD54 as a NEK1 target has been challenged recently (Ghosh *et al*, 2022).

Mutations in the human *NEK1* gene have been linked to several distinct disease aetiologies. For example, *NEK1* mutations have been found in human patients with autosomal recessive Majewski type short-rib polydactyly syndrome (SRPS), which is associated with polycystic kidneys (Chen *et al*, 2012; Thiel *et al*, 2011), lethal skeletal dysplasia, polydactyly, facial dysmorphism, drastic growth defects *in utero* and microcephaly (Chen *et al*., 2012). *NEK1* mutations have also been found in other skeletal dysplasias such as Jeune syndrome (McInerney-Leo *et al*, 2015) and axial spondylometaphyseal dyplasia (SMD) (Wang *et al*, 2017). The disease most strongly associated with *NEK1* mutations is amyotrophic lateral sclerosis (ALS), a form of motor neuron disease with symptoms that do not overlap with SMD or SRPS (Brenner & Freischmidt, 2022; Yao *et al*, 2021). In 2015, an exome sequencing-based report revealed heterozygous *NEK1* mutations in sporadic ALS (sALS) (Cirulli *et al*, 2015). Since then, a range of studies have validated *NEK1* as an ALS gene in both sALS and familial ALS (fALS) (Brenner *et al*, 2016; Goldstein *et al*, 2019; Gratten *et al*, 2017; Kenna *et al*, 2016; Nguyen *et al*, 2018; Riva *et al*, 2022; Shu *et al*, 2018; Tripolszki *et al*, 2019). The human NEK1 protein can be divided roughly into 3 regions: the N-terminal kinase domain, the central coiled-coil region implicated in protein-protein interaction and the C-terminal acidic region of unknown function (Fig. S1). The *NEK1* mutations associated with ALS, SMD and other diseases results in amino acid changes in all three regions of the protein, with little evidence for clustering. Most ALS-associated *NEK1* mutations are heterozygous and are presumed to be dominant and loss-of-function in nature but this remains to be determined (Goldstein *et al*., 2019; Kenna *et al*., 2016). At present, the impact of the pathological mutations on NEK1 kinase activity is not known, and it’s not understood how these mutations affect DNA repair, ciliogenesis or mitotic progression.

Mutations in the *C21ORF2* gene phenocopy *NEK1* mutations in Jeune syndrome, SMD and ALS (McInerney-Leo *et al*, 2017; van Rheenen *et al*, 2016; Wang *et al*, 2016); this pattern of similarity is unique, suggesting that the functions of these two genes are intimately linked. Consistent with this idea, C21ORF2 was identified by mass spectrometry of epitope tagged NEK1 stably expressed in human cells and vice versa; moreover C21ORF2, like NEK1, has been linked to control of primary cilia (Cirulli *et al*., 2015; Wheway *et al*, 2015) and homologous recombination (Fang *et al*, 2015). These studies led us to the hypothesis that C21ORF2 might be a regulatory subunit of NEK1, and that the interaction of NEK1 with C21ORF2 may be essential for NEK1 function in cells. To test these ideas, we set out to generate high quality antibodies and gene knockouts in untransformed cell lines to assess rigorously whether NEK1 and C21ORF2 form a complex at the endogenous level in human cells; to predict the structure of the interaction interface; to characterise the complex in detail and define interaction domains; to compare directly the phenotypic defects in NEK1- and C21ORF2-knockout cells; to establish a “rescue” system to carry out structure function analysis in phenotypic analyses. These experiments revealed that NEK1 kinase activity and interaction with C21ORF2 are essential for its function in cells, and that a subset of pathological mutations in NEK1 C-terminal acidic region abolish the interaction with C21ORF2.

## Results

### Generation of NEK1 and C21ORF2 antibodies and knockout cell lines

To enable us to rigorously test the interaction of endogenous NEK1 and C21ORF2 we generated polyclonal antibodies in sheep. As shown in Fig. 1A, affinity purified antibodies raised against the full length human C21ORF2 antibody recognised a band of approximately 28 kDa in extracts of untransformed ARPE-19 retinal pigmented epithelial cells, that was reduced in intensity by a C21ORF2-specific siRNA. Similarly, NEK1 antibodies recognised a band of the expected molecular weight (approximately 141 kDa), and band intensity was reduced by a NEK1-specific siRNA (Fig. 1A). Intriguingly, C21ORF2 levels were significantly reduced in NEK1-depleted cells suggesting they form a complex in cells.

**Figure 1.**
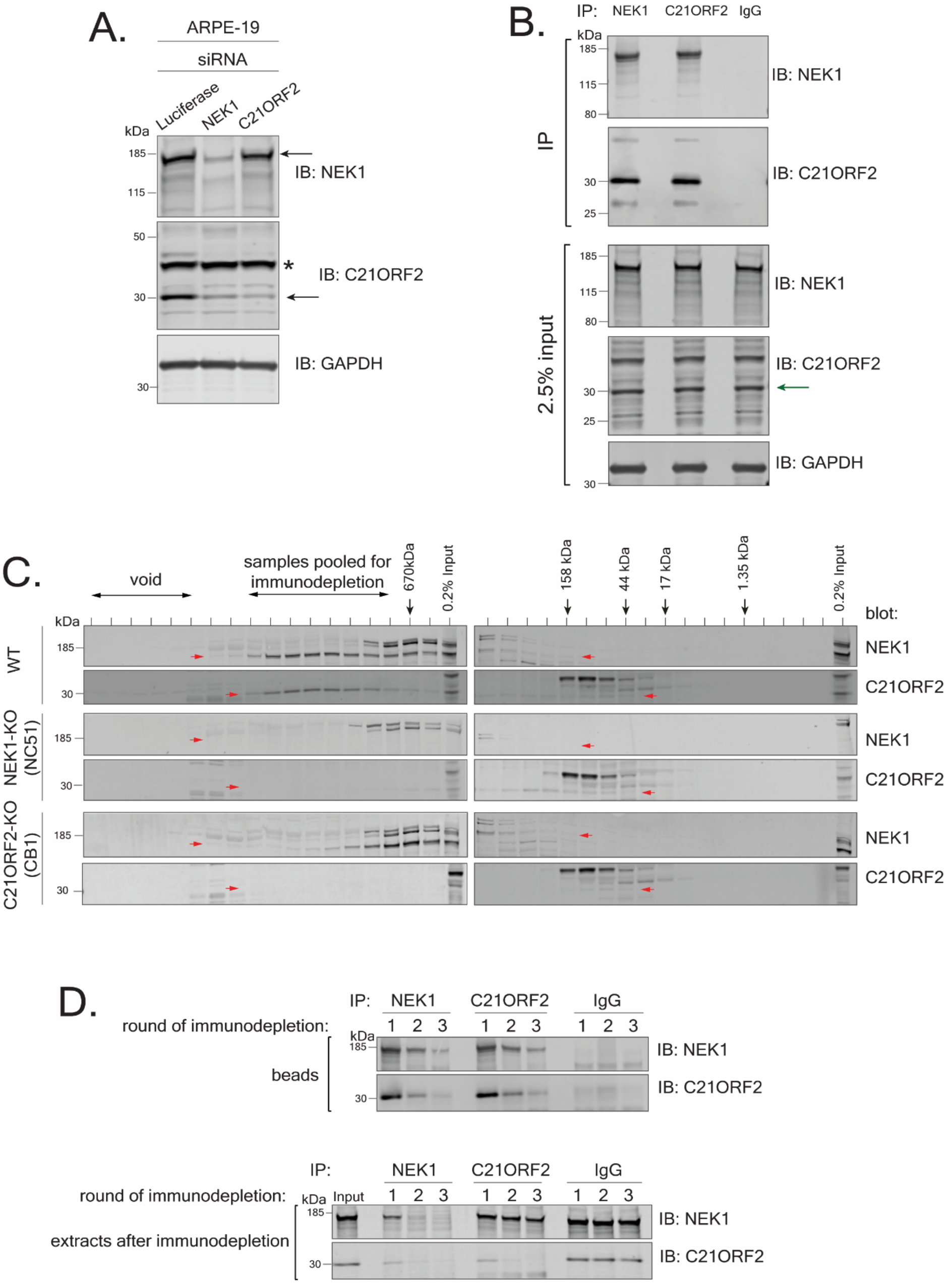
Characterization of the endogenous NEK1-C21ORF2 complex. (A) ARPE-19 cells were transfected with siRNA targeting either C21ORF2 or NEK1. Cell lysates were subjected to immunoblotting using in-house sheep polyclonal antibodies against C21ORF2 or NEK1 antibodies or GAPDH antibodies. Cell lysates treated with siRNA targeting luciferase served as a negative control. Asterisk denotes non-specific band; arrow denotes the specific C21ORF2 band. One of at least three independent experiments is shown. (B) ARPE-19 cells extracts were subjected to immunoprecipitation with in-house sheep antibodies against NEK1 or C21ORF2, or sheep IgG. Precipitates were subjected to SDS-PAGE and immunoblotting with the antibodies indicated, and input cell extracts were also included. Arrow denotes the C21ORF2 band. One of at least three independent experiments is shown. (C) Extracts from parental ARPE-19, NEK1-KO (clone NC51), and C21ORF2-KO (clone CB1) cells were subjected to size exclusion chromatography using a Superose 6 Increase 10/300 column. Alternate fractions were subjected to SDS–PAGE gel followed by immunoblotting using antibodies against NEK1 (Bethyl) and C21ORF2. The elution positions of molecular mass markers are indicated with black arrows. Red arrows indicate bands corresponding to NEK1 or C21ORF2. One of two independent experiments is shown. (D) The fractions indicated in C were pooled and subjected to three rounds of immunodepletion with either in-house antibodies against NEK1or C21ORF2 or sheep IgG antibodies. Beads and supernatants were analysed by SDS-PAGE and western blotting the antibodies indicated. Molecular weight markers “kDa” are indicated. One of two independent experiments is shown.

To further test the specificity of the antibodies, the *NEK1* and *C21ORF2* genes were disrupted in ARPE-19 cells. CRISPR-mediated genome editing was employed to disrupt *NEK1* using three distinct guide RNA (gRNA) pairs targeting exon 3 and 7. After screening over 100 single cell clones by western blotting, one clone was isolated which lacked any detectable NEK1 protein judged by blotting of extracts and immunoprecipitation using our NEK1 polyclonal antibodies (Figs. S2A, B; clone NC51). Sequencing of the genomic DNA near the gRNA target site in clone NC51 revealed that both alleles of *NEK1* had been disrupted. As shown in Fig. S2C, one allele had a large insertion at the end of exon 7 resulting in a premature STOP codon, while the other allele had a 10 bp deletion in exon 7, generating premature STOP codons at the beginning of exon 8. Clone NB32 was a hypomorph expressing very low levels of a truncated NEK1 protein (Figs. S2A, B; data not shown) and was not taken forward. As seen with NEK1 siRNA, C21ORF2 protein abundance was reduced in NEK1 clones NC51 and NB32 (Figs. S2A, B); C21ORF2 levels in clone NC51 were not rescued by proteasome inhibition (MG-132), inhibition of cullin NEDDylation (MLN-4294) or inhibition of autophagy (bafilomycin A or ULK1 inhibitor MRT68921) (Fig. S3). Therefore, the fate of C21ORF2 in the absence of NEK1 is not clear.

Next, *C21ORF2* was disrupted using CRISPR-mediated genome editing using two pairs of sgRNAs, each targeting sequences in exon 4. After screening around 100 clones, 7 clones (CA2, CA5, CA40, CA61, CB1, CB16, and CB17) were selected for further analysis. Clones CA2, CA5, CA40, CA61, CB1 lacked any detectable C21ORF2 protein judged by western blotting of extracts (Fig. S4A) and immunoprecipitation (Fig. S4B). Genotyping revealed that in clone CB1, one allele harbored two insertions in exon 4 and the other allele had simultaneous insertion and deletion, resulting in premature STOP codons in both alleles (Fig. S4C). Clone CB1 was taken forward for further analysis.

### Characterization of the endogenous NEK1-C21ORF2 complex

Having ascertained the specificity of our NEK1 and C21ORF2 antibodies, we used our polyclonal antibodies in immunoprecipitation experiments. As shown in Fig. 1B, a strong C21ORF2 band was detected in NEK1 precipitates from extracts of ARPE-19 cells and vice versa, and similar results were obtained in the cancer cell lines HeLa, HEK293 and U2OS (Fig. S5). The wash IP buffers contained 500 mM NaCl and Triton X-100 indicating the interaction is robust. Gel filtration of extracts of ARPE-19 cells revealed that NEK1 and C21ORF2 co-eluted in a broad peak before the 670 kDa marker (Fig. 1C). NEK1 eluted at a lower molecular mass in upon gel filtration of extracts of C21ORF2 knockout (KO) clone CB1, but not at the size expected for monomeric NEK1; C21ORF2 was only present at low levels in extracts of NEK1-KO cells and eluted at a much lower molecular mass than in parental ARPE-19 cells.

To determine the proportion of C21ORF2 that is bound to NEK1, and vice versa, the gel filtration fractions from parental cells containing NEK1 and C21ORF2 were pooled and subjected to immunodepletion. As shown in Fig. 1D, three rounds of depletion with NEK1 antibodies fully depleted all of the NEK1 from the pooled fractions; under these conditions, C21ORF2 was also fully depleted, indicating that all of the C21ORF2 was bound to NEK1. After three rounds of immunoprecipitation with C21ORF2 antibodies, C21ORF2 was fully depleted from the pooled fractions but under these conditions, NEK1 was not fully depleted however, indicating that not all of the NEK1 is in a complex with C21ORF2 (Fig. 1D).

Although the predicted mass of a NEK1-C21ORF2 heterodimer is approximately 169 kDa, the gel filtration experiments showed that NEK1 and C21ORF2 co-eluted in a complex with a molecular mass greater than 670 kDa (Fig. 1C). This discrepancy could be explained if the complex was fibrous in nature, or if multimerization occurred. Alternatively, there may be other proteins present in the complex. To find the full complement of NEK1 interactors, NEK1 was immunoprecipitated from ARPE-19 parental cells, and NEK1-KO cells were to control for non-specific binding. Five biological replicates were used per cell line, and after trypsinization of the precipitates, peptides were labelled with tandem mass tags (TMT) enabling quantitative mass spectrometric analysis of the 10 samples which were pooled and analysed in parallel. The experiment revealed that the only NEK1 interactor found to be at least twofold more abundant in NEK1 precipitates from parental cells compared with NEK1 KO cells was C21ORF2. Although, COL8A1 and C19ORF53 were also found to differ significantly between WT and NEK1 KO cells, they were treated as contaminants because the P-value was close to the 0.05 cut off (Fig. 2A). We also did the experiment the other way round, immunoprecipitating C21ORF2 from parental ARPE-19, with C21ORF2-KO cells used as control. The only C21ORF2 interactor which was found to be at least twofold more abundant in C21ORF2 precipitates from parental cells compared with C21ORF2-KO cells was NEK1 (Fig. 2B). These data reinforce the notion that C21ORF2 is the major NEK1 interactor, and argue that, at least under resting, asynchronous, unchallenged conditions there are no other major components of the endogenous NEK1-C21ORF2 complex.

**Figure 2.**
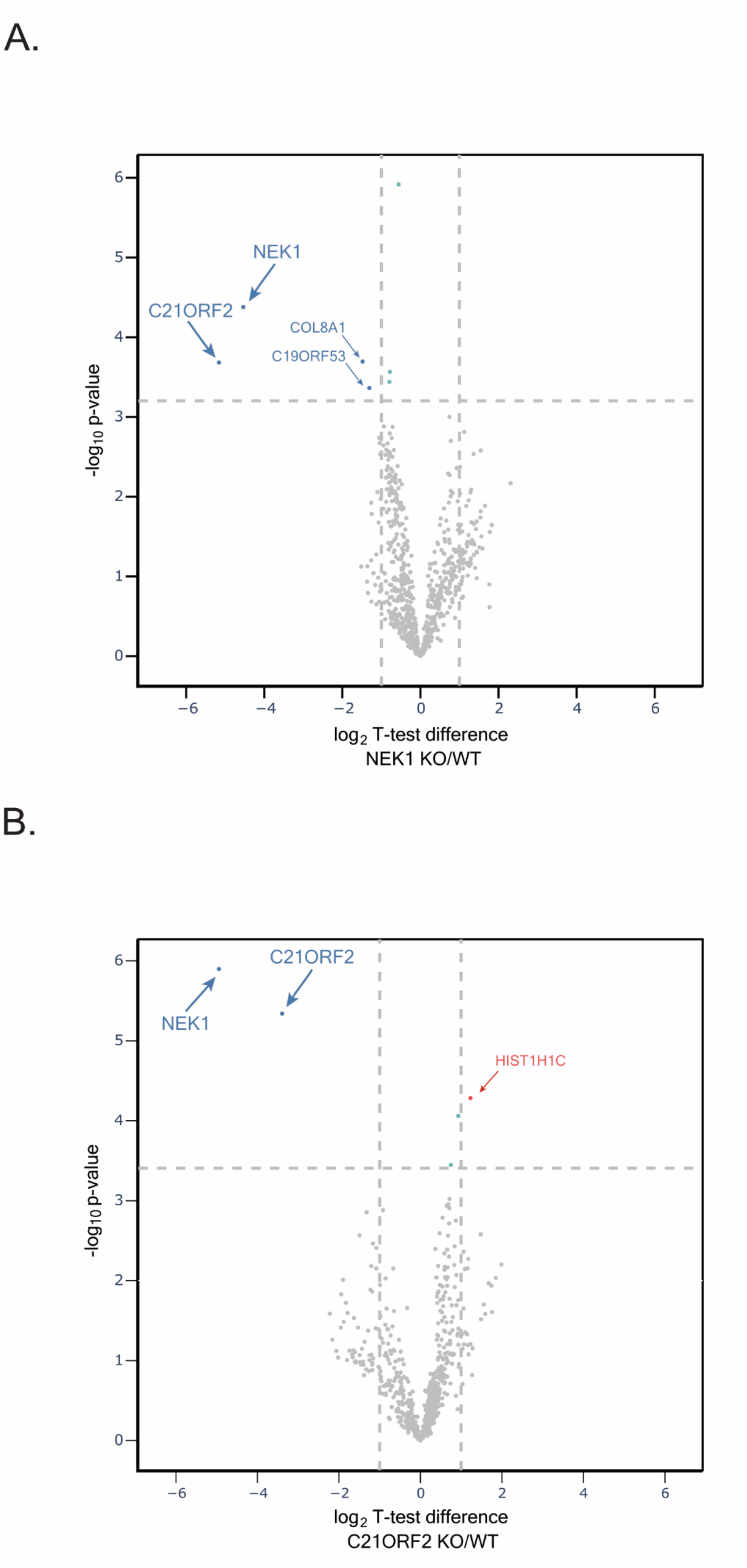
Mass spectrometric analysis of the NEK1–C21ORF2 complex in ARPE19 cells. (A) Lysates of ARPE-19 parental cells and NEK1–KO cells were subjected to immunoprecipitation with in-house sheep anti-NEK1 antibodies (5 biological replicates, 5mg per replicate). Proteins were eluted from beads, loaded on S-Trap columns, and after trypsinization, TMT-labelled samples were pooled and injected on an UltiMate 3000 RSLCnano System coupled to a Orbitrap Fusion Lumos Tribrid Mass Spectrometer. A volcano plot representing NEK1 interactors is shown. The horizontal cut off line represents a P-value of 0.05, and the vertical cut-off lines represent a log_2_ fold change above which peptides were considered to differ significantly in abundance between ARPE-19 WT and NEK1 KO cells. (B) Same as in A except extracts of ARPE-19 parental cells and C21ORF2–KO cells were subjected to immunoprecipitation with anti-C21ORF2 antibodies.

### Molecular determinants of the NEK1-C21ORF2 interaction

We next set out to determine the region of NEK1 interacting with C21ORF2, by investigating the ability of truncated versions of FLAG-tagged NEK1 to interact with GFP-tagged C21ORF2 in ARPE-19 cells (Fig. S6A). As shown in Fig. 3A, a fragment of NEK1 corresponding to the acidic domain at the C-terminus (aa acids 760-1286) co-precipitated with C21ORF2, similar to full length NEK1 whereas fragments corresponding to the kinase domain plus basic domain (aa 1-379), or the coiled coil domain (aa 379-760) did not. Similar results were obtained when GFP-C21ORF2 precipitates were probed for the FLAG-NEK1 fragments (data not shown). We tried to narrow down the C21ORF2 interacting domain further by making more NEK1 deletion constructs (Fig. S6A). These efforts showed that a small C-terminal fragment of NEK1 corresponding to aa 1160-1286 co-precipitated with C21ORF2, whereas deletion of this region prevented NEK1 from interacting with C21ORF2 (Fig. 3B). Thus, a small domain between residues 1160 and 1286 within the acidic region of NEK1 is necessary and sufficient to interact with C21ORF2 – we refer to this region as the NEK1-CID “C21ORF2-interacting domain”.

**Figure 3.**
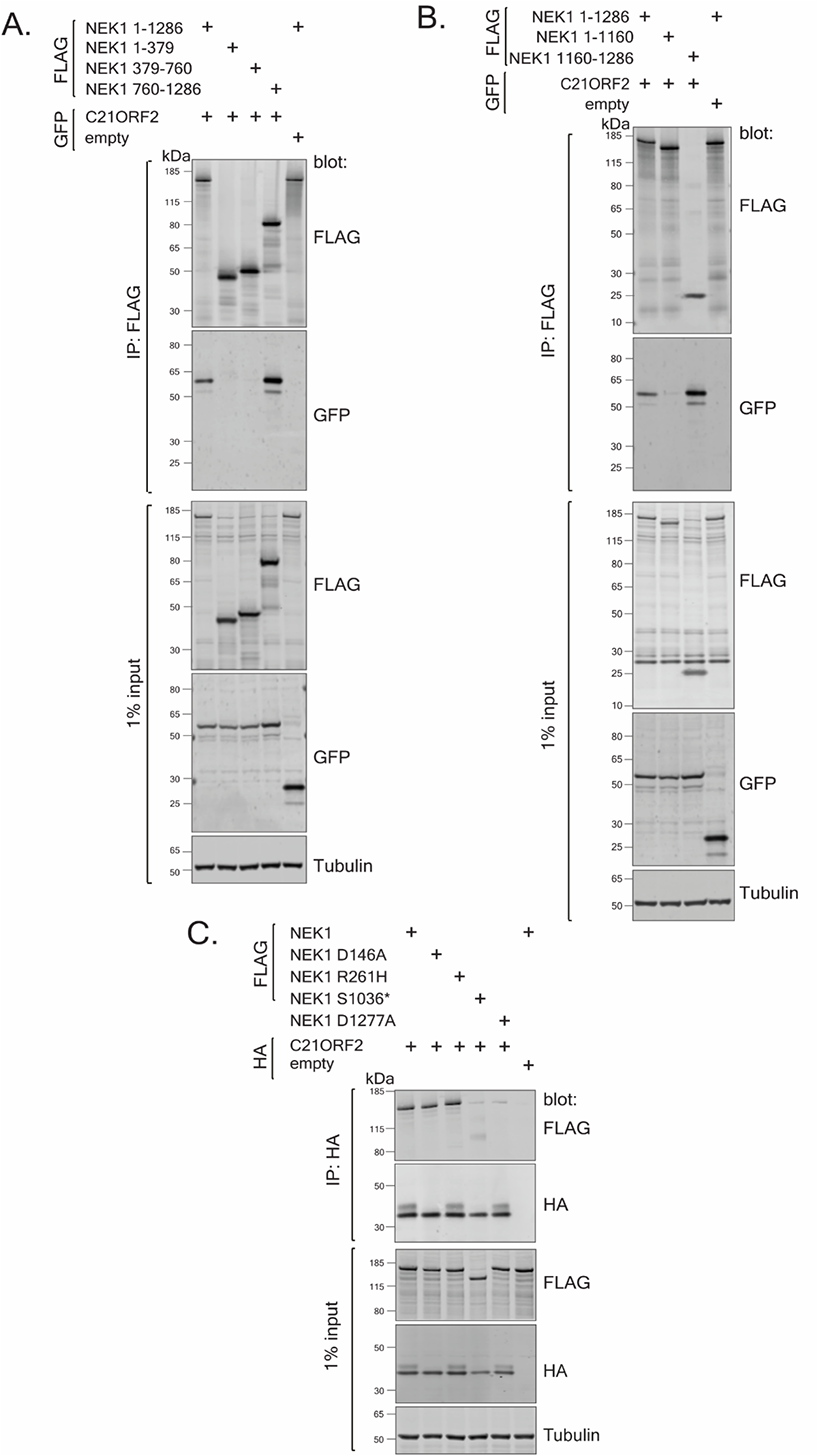
Molecular determinants of the NEK1-C21ORF2 interaction. (A, B) Lysates of ARPE-19 transiently co-transfected with plasmids encoding C21ORF2 (tagged with GFP on the N-terminus) and either full length or truncated forms of NEK1 (tagged with a 3xFLAG tag on the N-terminus) were subjected to immunoprecipitation with anti-FLAG or anti-GFP antibodies as indicated. Precipitates (and input cell extracts) were subjected to SDS-PAGE and immunoblotting with the indicated antibodies. One of two independent experiments is shown in each case. (C) Same as (A,B), except that ARPE-19 cells were co-transfected with cDNA encoding for C21ORF2 (tagged with 3xHA tag on the N-terminus) and wild-type or mutated versions of NEK1 (tagged with a 3xFLAG tag on the N-terminus). Molecular weight markers “kDa” are indicated. One of at least independent experiments is shown.

A small number of pathogenic mutations affect amino acids within or near the NEK1-CID defined above, raising the possibility that these mutations could disrupt the NEK1-C21ORF2 complex (Fig. S1). These include a mutation found in ALS and SMD which truncates NEK1 at S1036 (S1036*) (Kenna *et al*., 2016; Wang *et al*., 2017), and the D1277A mutation found in SMD (Wang *et al*., 2017) (Fig. S6A). Recent gene-burden analyses have revealed that R261H mutation in NEK1 is the most prevalent NEK1 ALS-associated mutation in the European patient cohort, and we also engineered this mutation in NEK1 (Kenna *et al*., 2016) (Fig. S6A). In order to test the impact of these mutations, ARPE-19 cells were transiently co-transfected with expression plasmids encoding N-terminal tagged 3HA-C21ORF2 and either wild-type or the NEK1 pathogenic variants described above (tagged with 3xFLAG on the N-terminus). As shown in Fig. 3C, 3xHA-C21ORF2 was detected in 3xFLAG-NEK1 precipitates, and the R261H NEK1 mutant had no detectable impact on binding to C21ORF2. However, the S1036* and D1277A NEK1 mutants showed a dramatic reduction in their interaction with C21ORF2, and the reverse co-immunoprecipitation experiment confirmed these findings (Fig. 3C). We also noted that a “kinase-dead” (KD) NEK1 mutant bearing a substitution in the ATP binding pocket (D146A) was indistinguishable from wild-type NEK1, suggesting that NEK1 kinase activity does not influence its binding to C21ORF2 (Fig. 3C).

The R73P and L224P amino acid substitutions in C21ORF2 found in ALS (van Rheenen *et al*., 2016) and Jeune Syndrome (Wheway *et al*., 2015), respectively, might abolish interaction with NEK1 (Fig. S6B). In order to test this possibility, ARPE-19 cells were co-transfected with plasmids encoding FLAG-NEK1 and either 3xHA-C21ORF2 (wild-type or mutants). Additionally, a C21ORF2 T150I mutant, recognised as the most prevalent C21ORF2 mutation in the European and US cohorts of ALS patients, was included (van Rheenen *et al*., 2016) (Fig. S6B). As shown in Fig. S6C, FLAG-NEK1 was detected in HA-C21ORF2 precipitates, but the R73P and L224P C21ORF2 substitutions weakened (but did not abolish) the interaction with NEK1. In contrast the C21ORF2 T150I mutant did not impact binding to NEK1; reverse co-immunoprecipitation experiment confirmed these findings (Fig. S6C). Taken together, the data above indicate that the NEK1-C21ORF2 interaction is disrupted by NEK1 pathogenic mutations S1036* and D1277A, and by the C21ORF2 R73P and L22P mutations.

### AlphaFold-based structural modelling of the NEK1-C21ORF2 interaction interface

We next used the ColabFold notebook (Mirdita *et al*, 2022) to run the AlphaFold structure prediction (Jumper *et al*, 2021) of the NEK1-C21ORF2 complex, using full length C21ORF2 and the NEK1-CID (aa 1160-1286) as input sequences. AMBER structure relaxation was used to ensure appropriate orientation of the side chains to avoid steric clashes. Five models were generated that were ranked from higher to lower confidence based on inter-chain predicted alignment error (inter-PAE) values (File S1) (Mirdita *et al*., 2022). All five models predicted a highly consistent binding interface between a region in the N-terminal half of C21ORF2 (aa 1-138) and a stretch of the NEK1-CID between residues 1216-1282 with high pLDDT and global PAE confidence scores for interacting regions (Figs. 4A, 7; S7 includes regions predicted to be disordered). This region of C21ORF2 appears to form a leucine-rich repeat (LRR) domain where 4 parallel beta strands contact a series of alpha helices in the NEK1-CID located between residues 1216 and 1282. Asp1277 of NEK1 is predicted to form a hydrogen bond with Asn24 of C21ORF2, providing an explanation for why substitution of Asp1277 for Ala (D1277A) disrupts the interaction with C21ORF2 (Fig. 3C). Arg73 of C21ORF2 lies in a loop connecting two LRR repeats in the NEK1 interaction interface (Figs. 4B). The kink introduced by the pathogenic R73P substitution likely disrupts the LRR domain mediating interaction with NEK1, which could explain why this substitution weakens binding of C21ORF2 to NEK1 (Figs. 4B).

**Figure 4.**
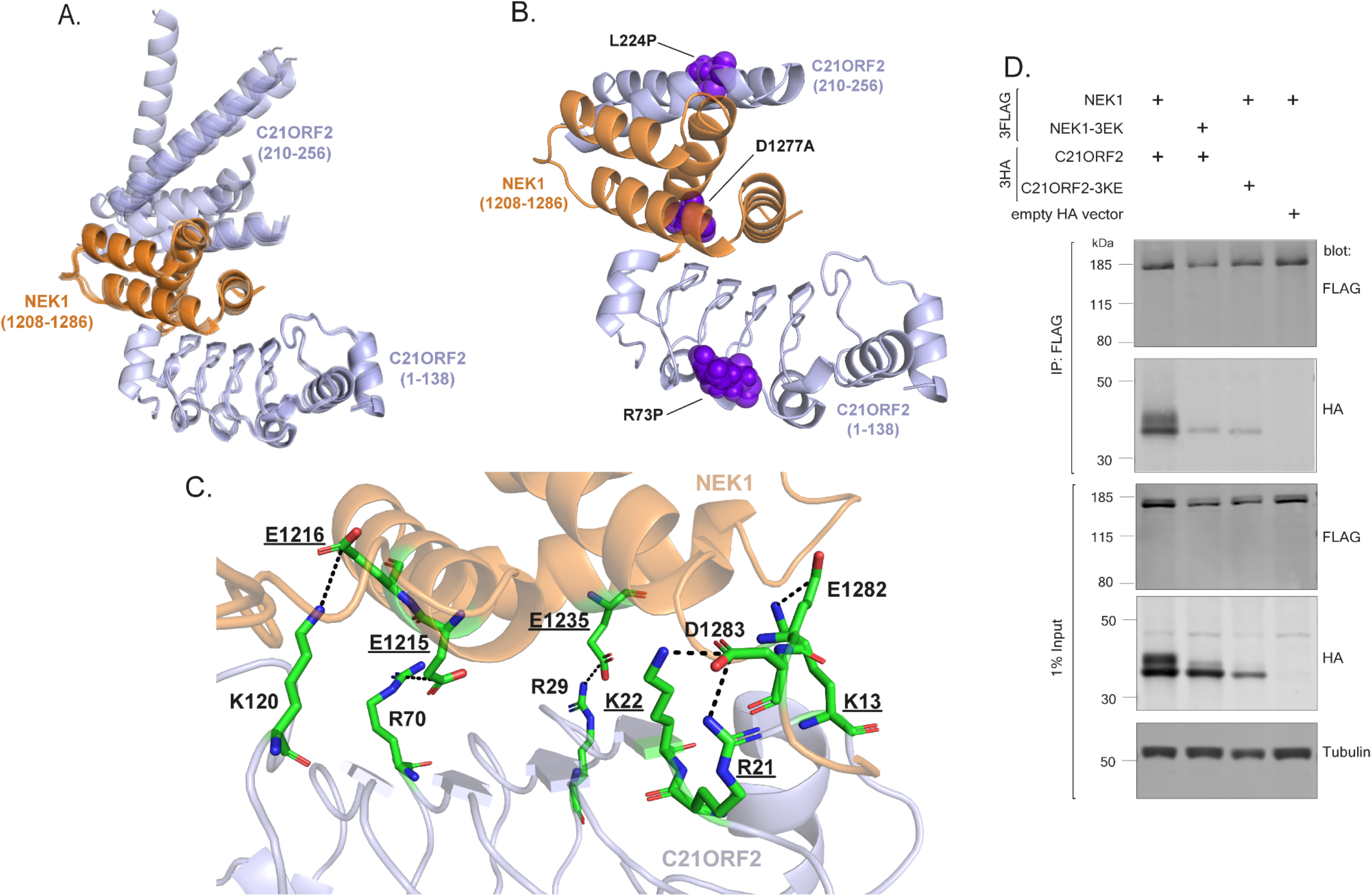
Alphafold structural modelling of the NEK1-C21ORF2 interaction interface. (A) Overlay of the 5 models of the NEK1–C21ORF2 interaction interface generated by Alphafold, with full length C21ORF2 and the NEK1-CID (aa 1160-1286) as the input sequences. The regions of C21ORF2 corresponding to residues 1-138 and residues 210-256 involved in NEK1 interaction are shown in light blue, while NEK1 residues 1208-1286 are shown in orange. (B) Amino acid substitutions encoded by pathological mutations in C21ORF2 [Arg73Pro (R73P) and Leu224Pro (L224P)] and NEK1 [Asp1277Ala (D1277A)], which disrupt the complex, map to the interaction interface predicted by the top ranked Alphafold model. The substitutions are marked in violet. The regions of C21ORF2 corresponding to residues 1-138 and residues 210-256 involved in NEK1 interaction are shown in light blue, while NEK1 residues 1208-1286 are shown in orange. (C) The binding interface between residues 1-138 in C21ORF2 (in orange) and NEK1 (in light blue) in the top ranked model highlighting amino acids that may mediate protein-protein interactions. Charged residues forming salt bridges between the proteins are marked (positively charged in blue, negatively charged in red). Residues selected for mutagenesis are underlined. (D) Lysates of ARPE-19 transiently co-transfected with plasmids encoding for C21ORF2 or C21ORF2-3KRE (tagged with 3XHA on the N-terminus) and NEK1 or NEK1-3EK (tagged with a 3xFLAG tag on the N-terminus). Precipitates and extracts were subjected western blotting with the antibodies indicated. One of two independent experiments is shown.

The interface between the N-terminal half of C21ORF2 (aa 1-138) and the NEK1-CID between residues 1216-1282 described above represents an extensive binding surface, with multiple contacts involving mostly electrostatic interactions and hydrogen bonds (Fig. 4C). A series of positively charged residues in C21ORF2 (Lys13, Arg21, Lys22, Arg29, Arg70, Lys120) are predicted to form salt bridges with a series of negatively charged residues within NEK1-CID (Glu1215, Glu1216, Glu1235, Glu1282, Asp1283) on a surface resembling a molecular Velcro (Fig. 4C). To test this binding model, we assessed the impact of C21ORF2 mutations (K13E+R21E+K22E; “3KRE” mutant) and NEK1 mutations (E1215K+E1216K+E1235K: “3EK” mutant) on partner binding. As shown in Fig. 4D, the NEK1-3EK and C21ORF2-3KE proteins show a pronounced reduction in ability to co-precipitate with partner protein, consistent with the Alphafold model above.

The top-ranked AlphaFold model also predicted a second binding interface involving the NEK1-CID helical core and a region at the C-terminus of C21ORF2 between residues 210-256 (Figs. 4A, B) albeit with low confidence. Here, two alpha helices of C21ORF2 are predicted to bind to the backside of the NEK1-CID. It is noteworthy that the pathogenic amino acid substitution L224P found in Jeune syndrome locates to the first of the two helices in the predicted interface, which may account for why this substitution weakens C21ORF2 binding to NEK1 (Fig. 4B). However, further structural analyses are needed to determine if this second interface is valid.

### C21ORF2-NEK1 is required for homologous recombination independent of RAD51

NEK1 has been implicated in a range of cellular functions, and we wondered if NEK1 interaction with C21ORF2 is important for these functions. With this in mind, we set out to compare C21ORF2-deficient cells with NEK1 deficient cells in a range of assays. NEK1 has been reported to control HR by phosphorylating Ser572 in the RAD54 ATPase (Spies *et al*., 2016). However, we have been unable to obtain evidence for RAD54 phosphorylation (data not shown). We examined RAD51 dynamics in NEK1- and C21ORF2-KO cells. As shown in Fig. 5A, after exposure of ARPE-19 cells to 2Gy IR, the proportion of cells with RAD51 foci peaked five hours post irradiation with around 60% of ARPE-19, NEK1 KO, and NEK1 KO + NEK1 cells containing five or more RAD51 foci. At this timepoint a slightly higher proportion (around 80%) of C21ORF2 KO and C21ORF2 KO + C21ORF2 cells had five or more RAD51 foci (Fig. 5A). Unloading of RAD51 was detected at the 10 h post irradiation, dropping to 40 % of parental ARPE-19 cells with greater than 5 RAD51 foci (Fig. 5A). There was no significant delay in unloading of RAD51 in *NEK1*-disrupted cells; although the proportion of cells with five or more RAD51 foci was higher in C21ORF2 KO, the dynamics of RAD51 unloading was comparable to parental cells. Our data suggest that NEK1 may not control the dynamics of RAD51 nucleofilament assembly or disassembly at IR-induced DSBs.

**Figure 5.**
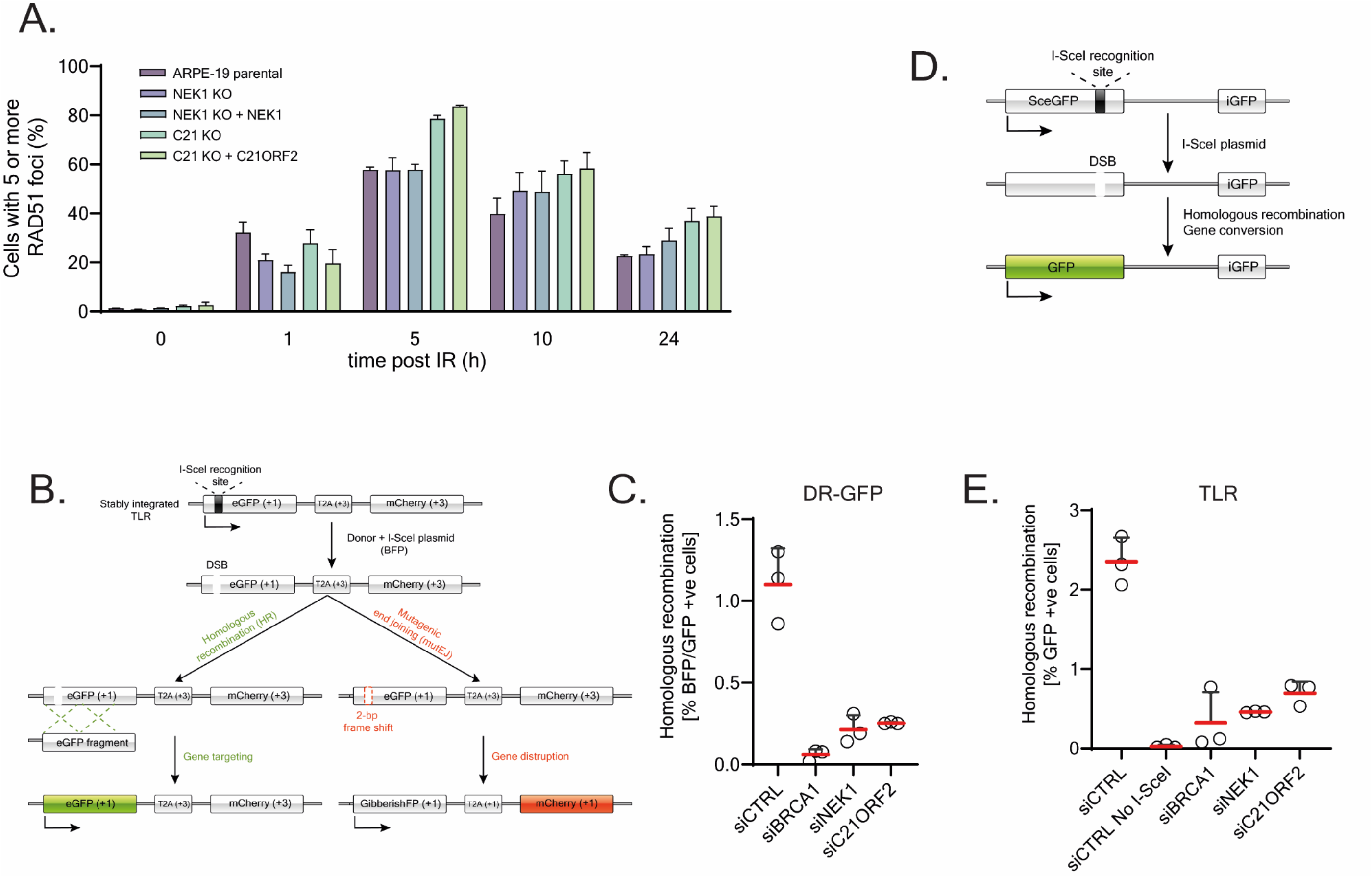
C21ORF2-NEK1 is required for homologous recombination independent of RAD51 dynamics. (A) ARPE-19 parental cells together with NEK1-KO, NEK1-KO + NEK1, C21ORF2-KO and C21ORF2-KO + C21ORF2 cells were subjected to an EdU pulse for 30 minutes followed by treatment with IR (2 Gy). Cells were allowed to recover for 1, 5, 10, and 24 h. Additionally, cells before IR treatment were collected to investigate the basal level of DNA damage. Cells were fixed, permeabilised, and stained for EdU and RAD51. DNA content was measured by staining cells with DAPI. Data was acquired with a high-content screening station ScanR; the proportion of EdU-positive cells with 5 or more RAD51 foci was counted. Data from three independent experiments was combined; data are represented as mean ± SEM. (B) Schematic diagram of TLR reporter. (C) U2OS TLR cells were transfected with siRNA and 48 h post transfection, cells were nucleofected with a bicistronic vector encoding GFP template and I-SceI nuclease. 24 h post plasmid transfection cells were harvested and analysed by flow cytometry for double positive BFP/GFP signals. Results from three independent experiments are shown. (D) Schematic diagram of DR-GFP reporter. (E) DR-GFP U-2OS cells were transfected with siRNA targeting either BRCA1, NEK1 or C21ORF2 or a non-targeting siRNA (siCTRL). 24 h later cells were nucleofected with a plasmid encoding I-SceI nuclease. 48 h post plasmid transfection cells were harvested and analysed by flow cytometry for GFP-positive cells. Results from three independent experiments are shown.

We next ascertained the involvement of NEK1 and C21ORF2 in HR by gene conversion and tested whether C21ORF2 is also required for this aspect of NEK1 function. We first used the “Traffic light reporter” (TLR) system (Certo *et al*, 2011) integrated into the genome of U2OS cells comprising a truncated *GFP* gene fused to an *mCherry* gene (Fig. 5B). The two genes are out of frame, blocking expression. Transfection of TLR cells with the endonuclease I-SceI generates a DSB in the *GFP* gene that, when repaired by gene conversion using an exogenous *GFP* template, leads to expression of a functional GFP (Fig. 5B). Transfection with a bicistronic plasmid encoding for an exogenous GFP template and I-SceI endonuclease can be monitored by detection of blue fluorescent protein (BFP). To test the role of NEK1 and C21ORF2 in HR, TLR cells were transfected with siRNA targeting NEK1 and C21ORF2, with BRCA1 siRNA or non-targeting siRNA serving as positive and negative controls, respectively. As shown in Fig. 5C around 1.1% of cells transfected with siCTRL underwent gene conversion after DSB induction. Depletion of NEK1 and C21ORF2 resulted a dramatic reduction in HR efficiency, similar in effect size to depletion of BRCA1.

We next used the well-characterised DR-GFP reporter system integrated into a U2OS genome which contains two *GFP* genes in a tandem arrangement (Fig. 5D) (Pierce *et al*, 1999). The upstream gene (*sceGFP*) is not functional as it includes a STOP codon within an I-SceI recognition site. The downstream gene (*iGFP*) is also inactive as it is truncated at both ends. Transient overexpression of I-SceI results in a DSB in the *SceGFP* which, if repaired by HR using the *iGFP* as a template, is restored to the fully functional *GFP* gene (Fig. 5D). DR-GFP cells transfected with siRNA targeting NEK1 or C21ORF2 were transfected later with a plasmid encoding I-SceI and analysed by flow cytometry 48 hours later. As shown in Fig. 5C, gene conversion at the I-SceI-induced DSB was observed in 2.3% of cells transfected I-SceI. As expected, no fluorescence was detected in siCTRL cells which were not transfected with I-SceI (Fig. 5E). Depletion of BRCA1 resulted in a 20-fold reduction in HR efficiency, and knockdown of NEK1 or C21ORF2 led to a comparable reduction (Fig. 5E). As HR only operates in S/G2 phases, we tested if the HR defects above could reflect altered cell cycle distribution. As shown in Fig. S8, depleting NEK1 caused a reduction in the relative proportion of S-phase cells, whereas depleting C21ORF2 had little effect. Given that the reduction in HR level after depletion of either NEK1 or C21ORF2 is similar in effect size, it is unlikely cell cycle distribution is responsible for the HR defect. Taken together, we conclude that cells deficient in NEK1 or C21ORF2 have a major defect in HR but that this function may not be related to control of RAD51 loading or unloading via RAD54 phosphorylation.

### A complementation system for structure-function-based phenotypic analyses

In the next phase of the study, we set out to test if C21ORF2-KO cells phenocopy NEK1-KO cells, and to test the functional impact of the pathogenic mutations which we found to disrupt the interaction of NEK1 and C21ORF2. Since the functional relevance of NEK1 kinase activity is unclear, we also wished to test the impact of a kinase-dead mutant. To enable these experiments, we established a system for re-expressing NEK1 or C21ORF2 in the respective knockout cell lines, that would not rely on protein overexpression. NEK1-KO cells were transduced with lentiviruses expressing NEK1 under the control of the CMV promoter which allowed expression at close to endogenous levels (Fig. 6A). A range of NEK1 mutants were also expressed: kinase-dead (KD) NEK1 (D146A), S1036* and D1277A. We noticed that both NEK1-WT and NEK1-KD restored C21ORF2 expression level back to normal in NEK1-KO cells (Fig. 6A). However, the C21ORF2 interaction defective NEK1 mutants did not rescue NEK1 expression, consistent with idea that complex formation regulates C21ORF2 stability (Fig. 6A). Consistent with transient overexpression-based experiments shown in Fig.3, the NEK1 D1277A mutant was unable to interact with C21ORF2 (Fig. 6A). We also transduced C21ORF2-KO cells with lentiviruses expressing C21ORF2 under the control of the UbC promoter which allowed expression at close to endogenous levels (Fig. 6B). We also introduced two pathogenic C21ORF2 mutants – L73P and L224P – found in ALS and Jeune Syndrome, respectively which only weakly interact with NEK1 (Fig. 6B).

**Figure 6.**
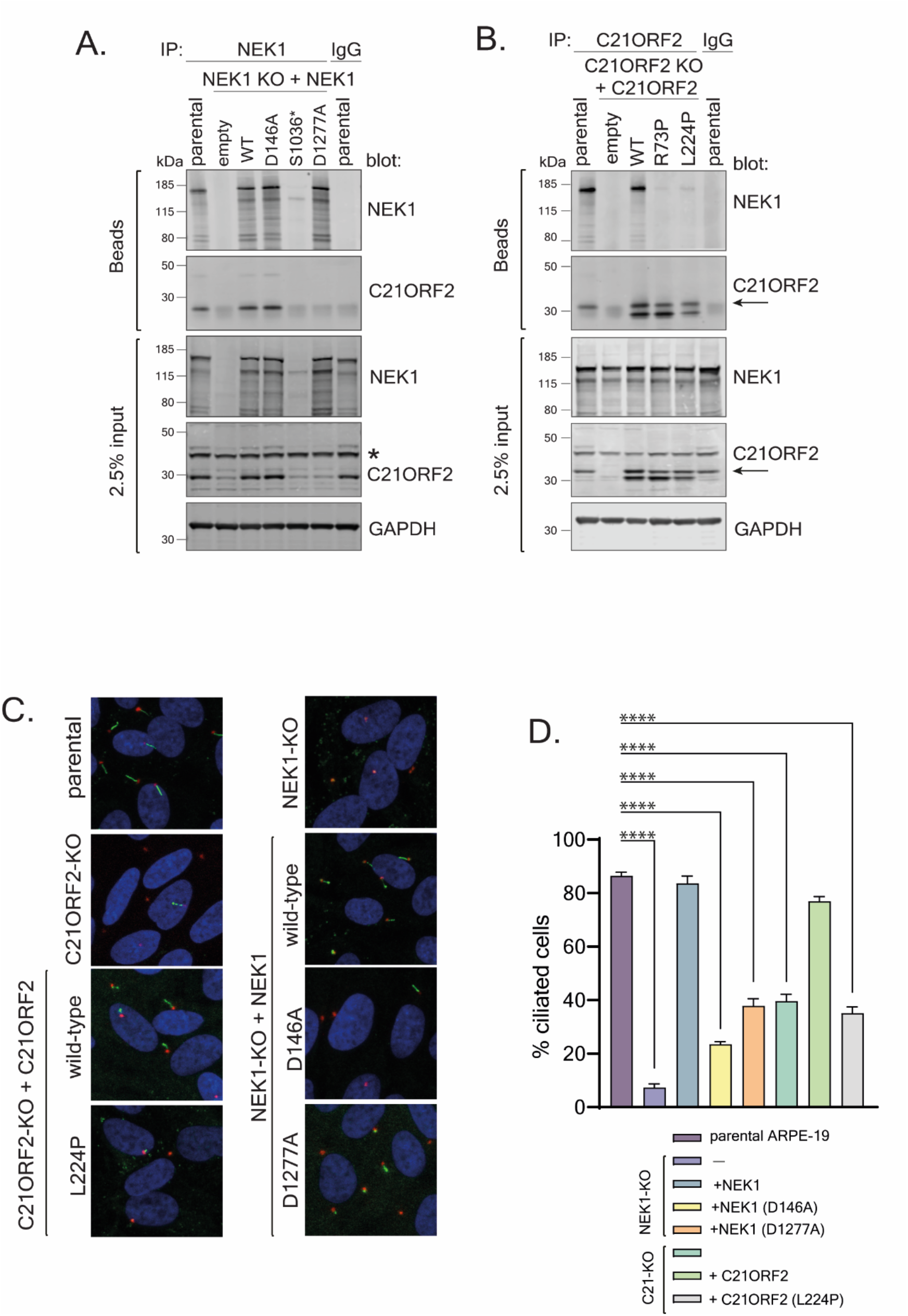
NEK1 kinase activity and interaction with C21ORF2 is critical for ciliogenesis. (A) NEK1 KO ARPE-19 cells (clone NC51) were transduced with lentivirus encoding wild-type (WT) NEK1 or the mutated versions of NEK1 indicated under the control of the CMV promoter; virus prepared from empty vector was used as control. Cells were selected for puromycin resistance, and extracts were subjected to immunoblotting with indicated antibodies; parental cell extracts were included. Extracts were also subjected to immunoprecipitation with antibodies against NEK1 and precipitates were blotted with the antibodies indicated. One of at least two independent experiments is shown. (B) C21ORF2 KO ARPE-19 cells (clone CB1) were transduced with lentivirus encoding wild-type (WT) C21ORF2 or the mutated versions of C21ORF2 indicated under the control of the UbC promoter; virus prepared from empty vector was used as control. Cells were selected for puromycin resistance, and extracts were subjected to immunoblotting with indicated antibodies; parental cell extracts were included. Extracts were also subjected to immunoprecipitation with antibodies against C21ORF2 and precipitates were blotted with the antibodies indicated. Molecular weight markers “kDa” are indicated. One of at least two independent experiments is shown. (C) ARPE-19 parental cells together with NEK1-KO cells, or NEK1-KO stably expressing NEK1, kinase-dead NEK1 (D146A) or NEK1 D1277A, C21ORF2-KO cells or C21ORF2-KO cells expressing C21ORF2 or C21ORF2 L224P were subjected to serum starvation for 48h followed by immunofluorescence with antibodies against ARL13B (green) or pericentrin (red). One of at least three independent experiments is shown. (D) Quantification of the proportion of ciliated cells in C. At least 150 cells were analysed for each condition per experiment. Data from three independent experiments was combined; data are represented as mean ± SD. Ordinary one-way ANOVA with multiple comparisons was used to evaluate the statistical significance of the results. p values: * p<0.05, ** p<0.01, *** p<0.001, **** p<0.0001.

### NEK1 kinase activity and interaction with C21ORF2 is critical for ciliogenesis

We next set out to test if NEK1 kinase activity and interaction with C21ORF2 are required for the formation of primary cilia. ARPE-19 cells were serum-starved for 48h to initiate exit to G_0_, and primary cilia were analysed by immunofluorescence using antibodies against ARL13B, an ARF-family small G protein highly enriched in cilia, and pericentrin to mark centrosomes. As shown in Figs. 6C and D, the proportion of NEK1-KO and C21ORF2-KO cells bearing primary cilia was reduced dramatically compared with parental cells. We noticed in some cells, that an ARL13B dot co-localized with a centrosome, suggesting that nucleation of microtubules may have started but microtubule extension had failed (Fig. 6C). Ciliogenesis was restored to normal by expression of NEK1 in NEK1-KO cells and expression of C21ORF2 in C21ORF2-KO cells (Figs. 6C, D). We also tested the impact of kinase activity and interaction with C21ORF2 on ciliogenesis. As shown in Figs. 6C and D, NEK1-KO cells expressing the kinase dead NEK1 D146A mutant or the D1277A mutant that cannot interact with C21ORF2 showed major defects in ciliogenesis. Therefore, NEK1 kinase activity and interaction with C21ORF2 are critical regulators of primary cilium formation.

## Discussion

In this study we have characterised the NEK1-C21ORF2 complex in human cells (Fig. 7). At least in APRE-19 cells, all of the C21ORF2 protein is bound to NEK1, but there is a pool of NEK1 that is not bound to C21ORF2. Given that the interaction with C21ORF2 of NEK1 appears to be essential for NEK1 function in ciliogenesis, and is likely to be required for HR, it is possible that the free pool of NEK1 is non–functional or has an alternative function. IP/MS analyses indicated NEK1 is the only major C21ORF2-interacting protein in resting cells, and *vice versa*, and no other complex components were detected (Fig 2). There is a report of the FLAG-tagged NEK1 overexpressed in HEK293T cells interacting with a variety of proteins after exposure of cells to cisplatin (Melo-Hanchuk *et al*, 2017). Although we did not detect these proteins in the NEK1-C21ORF2, they may bind to NEK1 after or C21ORF2 only after exposure of cells to genotoxins.

**Figure 7.**
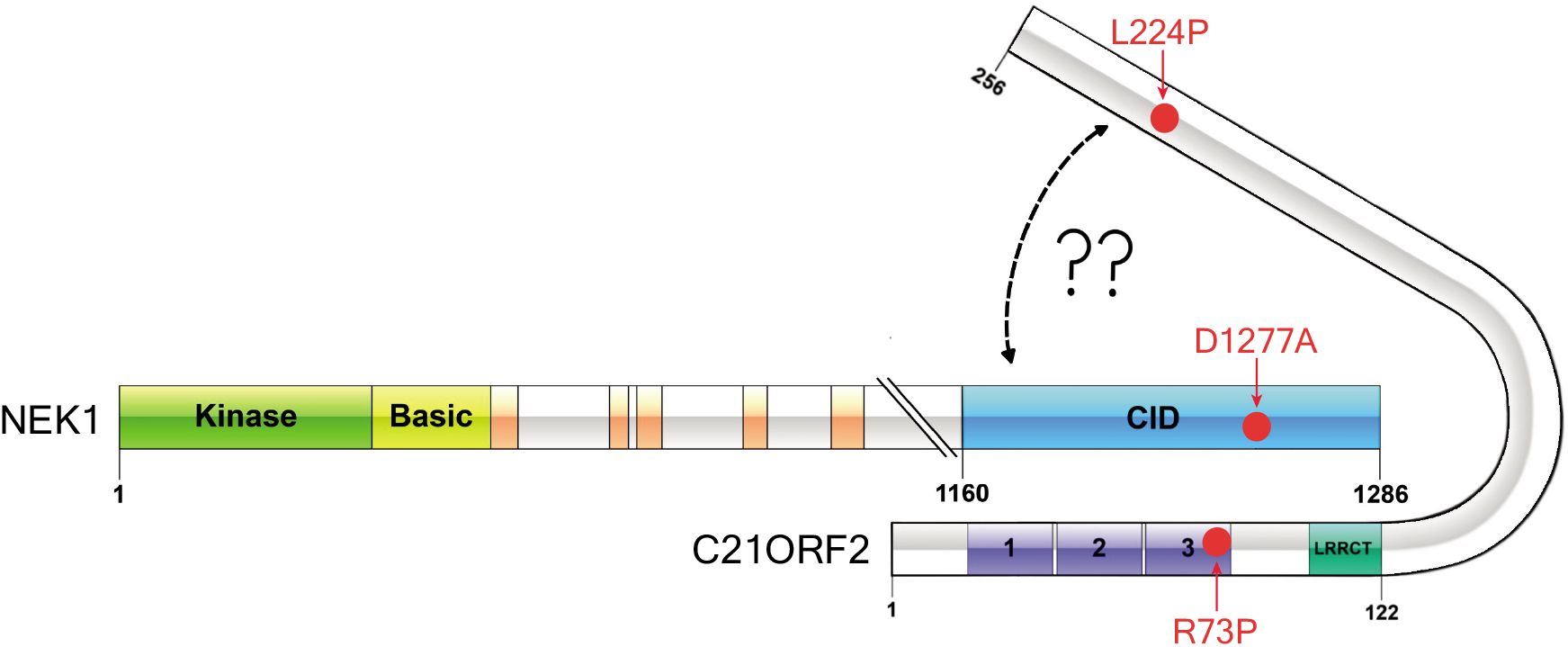
Schematic diagram of the NEK1-C21ORF2 complex. Globular domains in each protein are indicated. The NEK1-CID is highlighted in blue. AlphaFold modelling predicts with high confidence an interface between the N-terminal LRR-containing domain of C21ORF2 and the NEK1-CID. The top-ranked model predicts a second interface between the C21ORF2 C-terminal helices and the backside of the NEK1-CID but the confidence of this prediction is low and needs to validated.

We identified a small domain between amino acids 1160 and 1286 within the acidic, C-terminal region of NEK1, which we termed the CID, that is necessary and sufficient to interact with C21ORF2 (Fig. 7). AlphaFold modelling revealed an extensive interaction interface between the N-terminal LRR region of C21ORF2 and the NEK1-CID. This interface has an extended strip of positively charged residues lining the C21ORF2 side and a strip of negatively charged residues on NEK1 essential complex formation (Fig. 4). Importantly, the model explains why the ALS-associated D1277A NEK1 variant is unable to interact with C21ORF2, and defective in ciliogenesis (Fig. 7). A second interface between the C21ORF2 C-terminal helices and the backside of the CID was predicted with low confidence (Fig. 7); if valid, this model could potentially explain the pathogenicity of the L73P and L224P amino acid substitutions in C21ORF2 (Wheway *et al*., 2015). It will be important to validate the predictions made here, by solving the structure of the NEK1 C-terminus bound to C21ORF2, and it will be interesting to determine if the interaction in cells is regulated.

ARPE-19 cells lacking C21ORF2, like cells lacking NEK1, show major defects in ciliogenesis. Furthermore, mutations in NEK1 or C21ORF2 that disrupt the complex are unable to rescue the ciliogenesis defect in the respective knockout cells (Fig. 6). Therefore, the interaction mechanism defined in this study is critical for NEK1 function in cells. We also found that the kinase activity of NEK1 is essential for ciliogenesis (Fig. 6). The phenotypic similarity between the NEK1 kinase-dead mutant and the D1277A mutant unable to interact with C21ORF2 is striking. Taking all of these data together, perhaps the simplest model is that C21ORF2 is required for NEK1 to phosphorylate its substrates involved in ciliogenesis. Investigating this possibility is a top priority in terms of understanding the functional impact of C21ORF2 on NEK1, but at present the targets of NEK1 required for ciliogenesis are unknown. Generally speaking, we have not found a reliable biomarker we could use as a readout of endogenous NEK1 activity based on candidates from the literature, and no systematic phosphoproteomic screen for NEK1 targets has been reported. Therefore, identifying NEK1 targets is critical. Identifying the full complement of nuclear NEK1 targets will also be important given that we observed a major decrease in HR efficiency in cells depleted of NEK1, and in cells depleted of C21ORF2 in parallel (Fig. 5B-E). We have been unable to observe NEK1-dependent phosphorylation of RAD54 (data not shown), and in our hands RAD51 loading and unloading was unaffected in NEK1-KO or C21ORF2-KO cells after exposure to IR (Fig. 5A). More work will be needed to understand the regulation of HR by NEK1. It is interesting to note that the symptoms associated with SMD, caused by NEK1 or C21ORF2 mutations for example, are somewhat reminiscent of the SPONASTRIME syndrome caused by mutations in TONSL that weaken HR (Burrage *et al*, 2019; Chang *et al*, 2019). It is possible therefore, that SMD symptoms are caused by defective HR, and this idea will be important to test.

NEK1, like NIMA, is required for mitotic function and spindle integrity in mouse cells (Brieno-Enriquez *et al*., 2017; Chen *et al*., 2011) but it is not yet clear if this role requires interaction with C21ORF2. We set out to study mitotic progression in NEK1- and C21ORF2-KO human cells (or cells treated with siRNA) after release from a G_2_ arrest. However, we were unable to synchronise NEK1-deficient cells in G_2_ or M phases using a range of methods (data not shown). To rigorously test their roles in mitosis, we may need a way to switch off NEK1 and C21ORF2 acutely, using degron tags for example; these reagents are in the pipeline. It is tempting to speculate that the role of NEK1 in mitosis involves the phospho-dependent control of centrosomes and/or microtubules, and that this role is shared in ciliogenesis which, like spindle formation, requires plus-end microtubule extension from the centrosome. It is even possible that the role of NEK1-C21ORF2 in HR could be explained by microtubule regulation given reported links between the cytoskeleton, DSB mobility and DNA repair (Lottersberger *et al*, 2015). This will require further work and finding the substrates of NEK1-C21ORF2 in cells is a key priority that could provide valuable clues.

Overexpression of NEK1 but not the D146A kinase-dead mutant induced an electrophoretic mobility shift in co-expressed C21ORF2 (Fig. 3C) which is reversed by treating the C21ORF2 precipitates with lambda phosphatase. This suggests C21ORF2 is a NEK1 target, but we have never seen this shift in cells expressing endogenous levels of NEK1 (data shown). A recent report suggests that NEK1-mediated phosphorylation is necessary for C21ORF2 stability in overexpression-based experiments (Watanabe *et al*, 2020), but in our hands the reduced levels of C21ORF2 protein in NEK1-KO cells were restored by both wild-type and kinase-dead NEK1 (Fig. 6A). It will be important to map the phosphorylation sites in endogenous C21ORF2 and to investigate their functional significance.

Mutations in *C21ORF2* and *NEK1* have been associated with an overlapping set of distinct disease aetiologies, although in some cases it is not yet clear if the mutations are causal. The strongest link is to ALS, replicated across several independent studies, with *NEK1* now regarded as a *bona fide* ALS gene. However, it is not yet clear how the disease-associated mutations in NEK1-C21ORF2 affect the activity of the complex in cells at the molecular level. Do *NEK1* and *C21ORF2* mutations found in different diseases affect different aspects of NEK1-C21ORF2 function – ciliogenesis versus DNA repair, for example? The SMD symptoms caused by *NEK1* mutations are reminiscent of those reported in SPONASTRIME syndrome caused by mutations in the *TONSL* HR gene (Burrage *et al*., 2019; Chang *et al*., 2019). Therefore, it is possible that it is the HR function of NEK1 that is responsible for preventing SMD. On the other hand, genes involved in ciliary function have been implicated in preventing skeletal dysplasias and therefore it may be the ciliary defects associated with NEK1 that are the key to SMD (Chinipardaz *et al*, 2022). Do *NEK1* and *C21ORF2* mutations found in different diseases affect different subsets of NEK1 kinase targets; again, this will require identification and robust validation of NEK1 targets. Understanding the molecular defects associated with deficiency in the NEK-C21ORF2 complex may open avenues for new treatment for diseases caused by mutations in *NEK1* or *C21ORF2*.

## Material and Methods

**Table 1.** Plasmids used in this work

| Protein expressed | Plasmid | Catalogue number |
| --- | --- | --- |
| 3xFLAG-NEK1 | pcDNA5 FRT/TO-CMV | DU58176 |
| 3xFLAG-NEK1 D146A (KD) | pcDNA5 FRT/TO-CMV | DU63607 |
| 3xFLAG-NEK1 R261H | pcDNA5 FRT/TO-CMV | DU67947 |
| 3xFLAG-NEK1 S1036* (nt change C3107G) | pcDNA5 FRT/TO-CMV | DU63628 |
| 3xFLAG-NEK1 D1277A (nt change A3830C) | pcDNA5 FRT/TO-CMV | DU63629 |
| 3xFLAG-NEK1 1-379 | pcDNA5 FRT/TO-CMV | DU67494 |
| 3xFLAG-NEK1 379-760 | pcDNA5 FRT/TO-CMV | DU67594 |
| 3xFLAG-NEK1 760-1286 | pcDNA5 FRT/TO-CMV | DU67595 |
| 3xFLAG-NEK1 1-1160 | pcDNA5 FRT/TO-CMV | DU67593 |
| 3xFLAG-NEK1 1160-1286 | pcDNA5 FRT/TO-CMV | DU68531 |
| 3xHA-C21ORF2 | pcDNA5 FRT/TO-UbC | DU70174 |
| 3xHA-C21ORF2 R73P | pcDNA5 FRT/TO-UbC | DU70225 |
| 3xHA-C21ORF2 T150I | pcDNA5 FRT/TO-UbC | DU70230 |
| 3xHA-C21ORF2 L224P | pcDNA5 FRT/TO-UbC | DU70247 |
| 3xHA-empty | pcDNA5 FRT/TO-UbC | DU70173 |
| GFP-NEK1 | pcDNA5 FRT/TO-CMV | DU58446 |
| GFP-NEK2 | pcDNA5 FRT/TO-CMV | DU58431 |
| GFP-NEK3 | pcDNA5 FRT/TO-CMV | DU58453 |
| GFP-NEK4 | pcDNA5 FRT/TO-CMV | DU58432 |
| GFP-NEK5 | pcDNA5 FRT/TO-CMV | DU58433 |
| GFP-NEK6 | pcDNA5 FRT/TO-CMV | DU58432 |
| GFP-NEK7 | pcDNA5 FRT/TO-CMV | DU58440 |
| GFP-NEK8 | pcDNA5 FRT/TO-CMV | DU58443 |
| GFP-NEK9 | pcDNA5 FRT/TO-CMV | DU58444 |
| GFP-NEK10 | pcDNA5 FRT/TO-CMV | DU58430 |
| GFP-NEK11 | pcDNA5 FRT/TO-CMV | DU58895 |
| GFP-C21ORF2 | pcDNA5 FRT/TO-CMV | DU67626 |
| GFP-empty | pcDNA5 FRT/TO-CMV | DU41455 |
| NEK1 | pLV(exp)-puro-CMV | DU70683 |
| NEK1 D146A (KD) | pLV(exp)-puro-CMV | DU61644 |
| NEK1 S1036* | pLV(exp)-puro-CMV | DU61594 |
| NEK1 D1277A | pLV(exp)-puro-CMV | DU61595 |
| CMV empty | pLV(exp)-puro-CMV | DU70925 |
| C21ORF2 | pLV(exp)-puro-UbC | DU70547 |
| C21ORF2 R73P | pLV(exp)-puro-UbC | DU70719 |
| C21ORF2 L224P | pLV(exp)-puro-UbC | DU70720 |
| UbC-empty | pLV(exp)-puro-UbC | DU70930 |
| NEK1 KO NA Sense | pBabeD P U6 | DU57080 |
| NEK1 KO NA Antisense | pX335 | DU57088 |
| NEK1 KO NB Sense | pBabeD P U6 | DU57081 |
| NEK1 KO NB Antisense | pX335 | DU57089 |
| NEK1 KO NC Sense | pBabeD P U6 | DU57080 |
| NEK1 KO NC Antisense | pX335 | DU57088 |
| C21ORF2 KO CA Sense | pBabeD P U6 | DU64911 |
| C21ORF2 KO CA Antisense | pX335 | DU64913 |
| C21ORF2 KO CB Sense | pBabeD P U6 | DU64912 |
| C21ORF2 KO CB Antisense | pX335 | DU64914 |

**Table 2.**
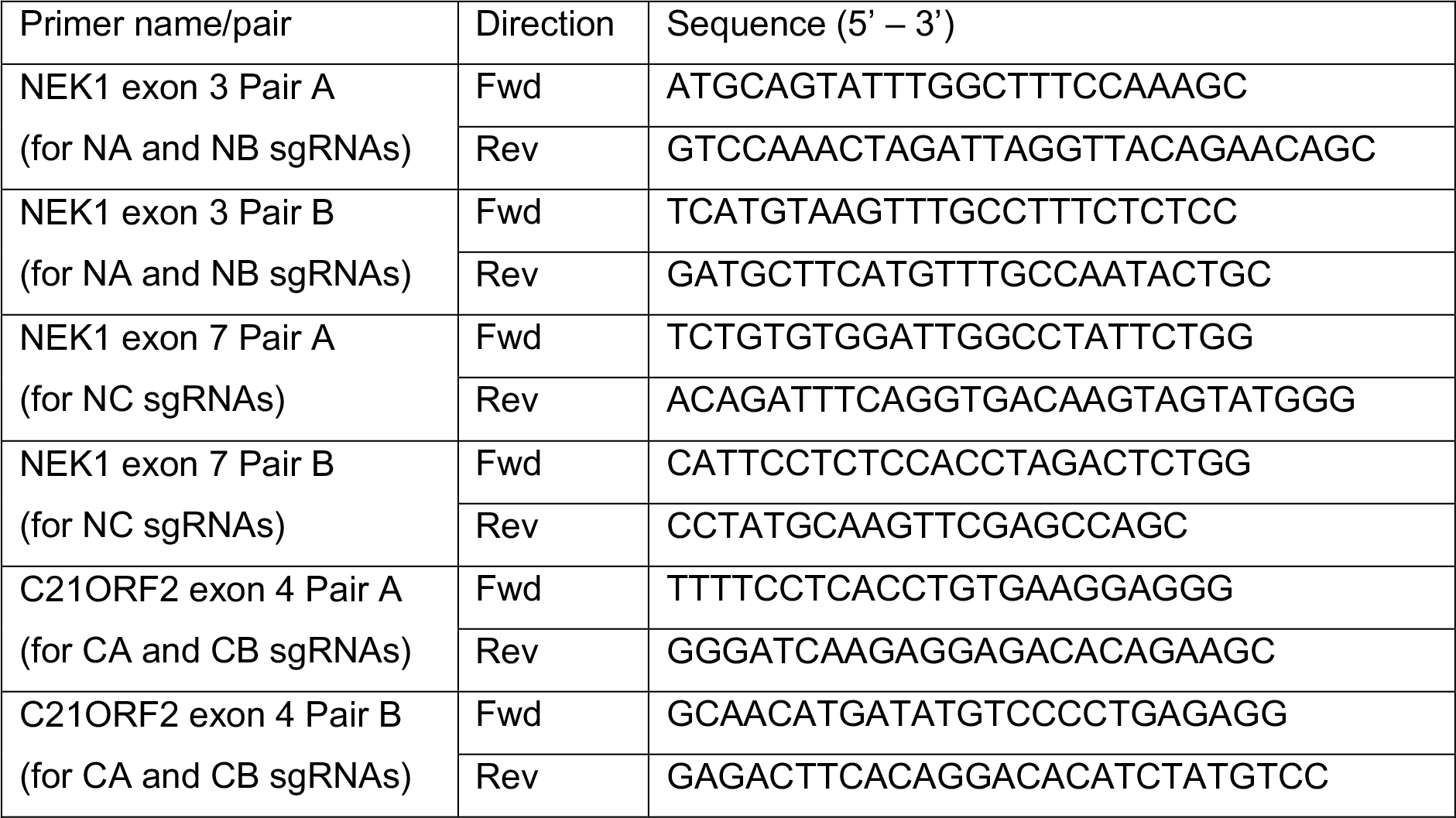
Oligonucleotides used in this work

**Table 3.**
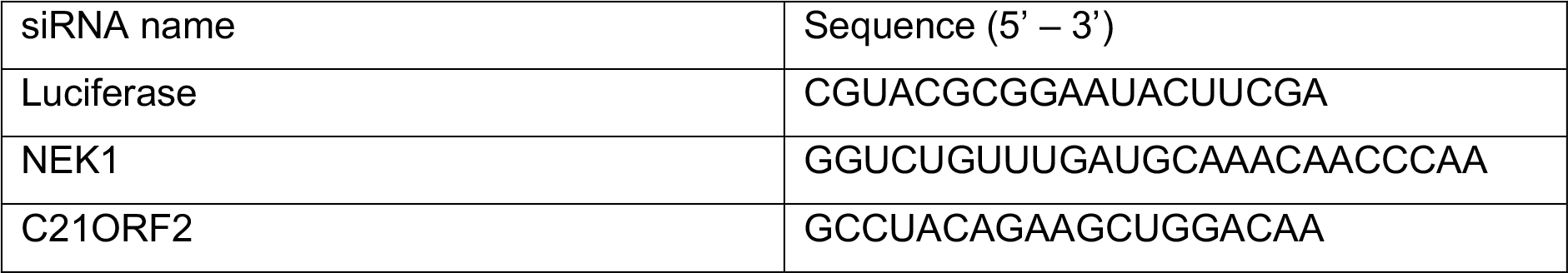
siRNAs used in this work

### Antibody production

Polyclonal NEK1 and C21ORF2 antibodies were raised in sheep by MRC-PPU Reagents and Services (University of Dundee) and purified against the relevant antigen (after depleting antibodies recognizing the epitope tags). NEK1: sheep SA354; 3rd bleed; antigen corresponded to GST-NEK1 (NM_001199397.1) aa 900-1286 expressed in bacteria. C21ORF2: sheep DA066, 4^th^ bleed; antigen corresponding to maltose binding protein-(MBP) tagged C21ORF2 aa 1-256 (NM_004928.2) expressed in bacteria). Sheep were immunised with the antigens followed by four further injections 28 days apart. Bleeds were performed seven days after each injection.

### Cell lines

All cell lines used in this study were derived from ARPE-19 cells. Cells were incubated at 37°C, 5% CO2 and maintained in Dulbecco’s Modified Eagle Medium/Nutrient Mixture F-12 (DMEM/F-12, ThermoFisherScientific) supplemented with 10% FBS (ThermoFisherScientific), 1X 10000 U/mL Penicillin-Streptomycin (ThermoFisher Scientific), 1/100 200 nM L-glutamate (ThermoFisher Scientific). ARPE-19 NEK1 KO and C21ORF2 KO cell lines were maintained as above with a culture medium supplemented with 20% FBS. Stably transduced ARPE-19 NEK1 KO and C21ORF2 KO cells were maintained as above with a culture medium supplemented with 20% FBS and 2µg/ml puromycin (ThermoFisherScientific). All cell lines were cultured in medium supplemented with 10% FBS during the duration of experiments. U2OS cells DR-GFP cells and U2OS TLR cells were cultured in DMEM medium supplemented with 10% FBS (Wisent, St-Bruno, Canada) and 1% penicillin/streptomycin (ThermoFisher Scientific).

### siRNA transfection

150,000 ARPE-19 cells were seeded in 5cm plate and allowed to recover for 24 hours. Cells were transfected with 50nM siRNA using Lipofectamine RNAiMAX (ThermoFisherScientific) according to the manufacturer’s instructions. 8 hours post transfection, cells were washed once with PBS and added fresh complete DMEM F/12 culture medium. Cells were harvested 64 hours later.

### PEI transfection

For transfection of one 70% confluent ARPE-19 10cm dish a mixture of 10µg cDNA, 20µl 1mg/ml PEI MAX (Polysciences) topped up to 1ml with OptiMEM medium was prepared. Transfection reaction was incubated 20 minutes at room temperature before being added to cells in a dropwise fashion. 8 hours post transfection, cells were washed once with PBS and added fresh complete DMEM F/12 culture medium. Cells were harvested 24 hours post transfection.

### Genome editing

ARPE-19 cells were transfected with a pair of plasmids targeting exon 3 or 4 in *NEK1* sequence or exon 4 in *C21ORF2* sequence using Lipofectamine 2000 (ThermoFisherScientific, #11668027) according to manufacturer’s instructions. 8 hours post transfection, cells were washed once with PBS and added fresh complete DMEM F/12 culture medium. Cells were allowed to recover for additional 40 hours before being selected for antibiotic resistance using complete DMEM/F12 medium supplemented with 2μg/ml puromycin. The puromycin media was refreshed after 24 h for a total selection time of 48 hours. Cells were then cultured for an additional 3–5 days to provide time for gene editing and eventually seeded at low densities (500 cells) in 15 cm dishes. Single colonies were isolated using cloning discs (Sigma) soaked with trypsin two to four weeks later.

### Retrovirus production for stable expression of target proteins

To reintroduce expression of WT or mutated versions of NEK1/C21ORF2, NEK1 or C21ORF2 KO cells were infected with lentiviruses. HEK293FT cells were transfected with plasmids encoding for proteins of interest along with the GAG/Pol and VSVG constructs required for lentiviral production. PEI was used as a transfection reagent. 48 hours later lentiviral-containing medium was collected, filtered through a 0.22 µm pore filter (Millipore) and supplemented with polybrene (8 µg/ml). Cells were transduced for 24 hours followed by a 48-hour selection with 2µg/ml puromycin. Successful lentiviral integration was confirmed though Western blotting.

### Cell lysis and immunoprecipitation

Cells were lysed in ice cold buffer comprising of 50 mM Tris-HCl (pH=7.4), 150mM NaCl, 270 mM sucrose, 1 mM EGTA (pH 8.0, Sigma), 10 mM β-Glycerol phosphate disodium salt pentahydrate (Sigma), 5mM sodium pyrophosphate decahydrate (Sigma), 50 mM sodium fluoride (Sigma), 10 ng/ml Microcystin-LR (Sigma), 0.5 U/ml Pierce Universal Nuclease (ThermoFisherScientific), 1:100 Phosphatase Inhibitor Cocktail 2 (Sigma), 1x Complete (EDTA free) protease inhibitor (Roche), 1mM sodium orthovanadate (Sigma), 1mM AEBSF, 1mM Benzamidine, 10mM Iodoacetamide (Sigma), and 1% (v/v) Triton-X100 (Sigma). Lysates when incubated on ice for 20 minutes followed by centrifugation at 14,000 g, 4°C for 15 min. Proteins were quantified using the BCA Kit (ThermoFisher Scientific). Firstly, lysates were pre-cleared with beads conjugated to a primary antibody isotype control for 1 h at 4°C with applied shaking (1400 rpm). Lysates were then transferred to tubes containing beads conjugated to primary antibodies. Reactions were incubated for 1.5 h at 4°C with applied shaking (1400 rpm). Protein-bound beads were washed 5 x 15 minutes at 4°C on a rotating wheel with a buffer composed of 50 mM Tris (pH=7.4), 500 mM NaCl and 1% Triton X-100 before being analysed by SDS-PAGE and Western Blotting. The in-house MRC-PPU-generated NEK1 and C21ORF2 were used in all relevant immunoprecipitation experiments.

### SDS-PAGE and Western Blotting

The proteins were separated by SDS-PAGE using 4-12% Bis/Tris gels (ThermoFisherScientific) under reducing conditions for 3 hours at 90V. NuPAGE 3-[N-morpholino] propane sulphonic acid (MOPS) running buffer was used (ThermoFisherScientific). Proteins were transferred to 0.45μM nitrocellulose membrane (Cyvita) at a constant voltage of 90V for 1.5 hours. Membranes were blocked for 1 hour at room temperature using 5% milk/TBST. Detection of antigens was carried out using antibodies specified in Table 4. All antibodies were dissolved in 5% milk/TBST and incubated overnight (at least 16 hours) at 4°C. Subsequently, membranes were washed 3×5 min using TBST. Primary antibodies were detected using secondary antibodies specified in Table 4. All secondary antibodies were dissolved in 5% milk/TBST and incubated for 1 hr at room temperature. Membranes were washed 3 x 10 min using TBST and 1×10 minutes using PBS. Data was acquired using Odyssey CLx LI-COR scanner and analysed in Image Studio v. 5.2. The in-house MRC-PPU-generated NEK1 and C21ORF2 were used in all western blotting experiments except for blotting gel filtration fractions in Fig. 1C where Bethyl NEK1 antibodies were used (see Table 4 above).

**Table 4.** Antibodies used in this work

| Primary antibodies |  |  |  |  |  |
| --- | --- | --- | --- | --- | --- |
| Antigen | Species | Manufacturer | Identifier | Dilution | Application |
| FLAG | Mouse | Sigma | F1804 | 1:5000 | WB |
| HA | Rabbit | CST | 3724 | 1:1000 | WB |
| NEK1 | Rabbit | ProteinTech | 27146-1-AP | 1:1000 | WB |
| NEK1 (aa 700-1286) | Sheep | MRC PPU | SA353 | 1:2000 | WB |
| NEK1 (aa 900-1286) | Sheep | MRC PPU | SA354 | 1:2000 | WB |
| NEK1 | Rabbit | Bethyl | A304-570A | 1:2000 | WB |
| C21ORF2 (full length) | Sheep | MRC PPU | DA066 | 0.1µg/ml | WB |
| C21ORF2 (full length) | Rabbit | ProteinTech | 27609-1-AP | 1:1000 | WB |
| GAPDH | Rabbit | CST | 2118 | 1:10000 | WB |
| α-Tubulin | Mouse | CST | 3873 | 1:10000 | WB |
| GFP | Chicken | Abcam | Ab13970 | 1:1000 | WB |
| HIF-1α | Mouse | BD | 610959 | 1:1000 | WB |
| ATG14 | Rabbit | CST | 96752 | 1:1000 | WB |
| ATG14 pSer29 | Rabbit | CST | 92340 | 1:1000 | WB |
| LC3A/B | Rabbit | CST | 4108 | 1:1000 | WB |
| FLAG | Mouse | Sigma | F1804 | 1µg/1mg lysate | IP |
| Mouse IgG1 (Isotype control for FLAG antibody) | Mouse | (CST) | 5415 | 1µg/1mg lysate | IP |
| HA beads | Mouse | MRC PPU | Frankenbody | 15µl 50% slurry/ 1mg lysate | IP |
| Mouse IgG2b (Isotype control for HA beads) | Mouse | CST | 53484 | 1µg/1mg lysate | IP |
| NEK1 | Rabbit | ProteinTech | 27146-1-AP | 2µg/1mg lysate | IP |
| NEK1 (aa 900-1286 aa) | Sheep 3<sup>rd</sup> bleed | MRC PPU | SA354 | 1µg/1ml lysate | IP |
| NEK1 | Rabbit | Bethyl | A304-570A | 2µg/1mg lysate | IP |
| C21ORF2 (full length) | Sheep 4<sup>th</sup> bleed | MRC PPU | DA066 | 1µg /1mg lysate | IP |
| C21ORF2 (full length) | Rabbit | ProteinTech | 27609-1-AP | 2µg/1mg lysate | IP |
| Rabbit IgG | Rabbit | CST | 2729 | 2µg/1mg lysate | IP |
| Sheep IgG | Sheep | Abcam | ab37385 | 1µg/mg lysate | IP |
| NEK1 (aa 900-1286) | Sheep | MRC PPU | SA344 | 1:100 | IF |
| C21ORF2<br>(full length) | Sheep | MRC PPU | DA066 | 1:200 | IF |
| GTU88 | Mouse | Sigma Aldrich | T6557 | 1:500 | IF |
| RAD51 | Rabbit | BioAcademia | 70-002 | 1:1000 | IF |
| γH2AX | Mouse | Merck Millipore | 05-636 | 1:2000 | IF |
| ARL13B | Rabbit | ProteinTech | 17711-1-AP | 1:200 | IF |
| Pericentrin | Mouse | Abcam | ab28144 | 1:500 | IF |
| Secondary antibodies |  |  |  |  |  |
| Antigen | Species | Manufacturer | Code | Dilution | Application |
| Anti-mouse<br>680 | Donkey | LI-COR | 926-68072 | 1:10000 | WB |
| Anti-mouse<br>800 | Donkey | LI-COR | 926-32212 | 1:10000 | WB |
| Anti-rabbit<br>680 | Donkey | LI-COR | 926-68073 | 1:10000 | WB |
| Anti-rabbit<br>800 | Donkey | LI-COR | 926-32213 | 1:10000 | WB |
| Anti-sheep<br>AF680 | Donkey | ThermoFisherScientific | A21102 | 1:10000 | WB |
| Anti-sheep<br>Dylight<br>800 | Donkey | Rockland | 613-745-168 | 1:10000 | WB |
| Anti-chicken<br>800 | Donkey | LI-COR | 926-32218 | 1:10000 | WB |
| Protein G<br>Dylight<br>800 | n/a | Rockland | PG00-45 | 1:10000 | WB |
| Anti-mouse<br>AF488 | Donkey | ThermoFisherScientific | A21202 | 1:1000 | IF |
| Anti-mouse<br>AF647 | Donkey | ThermoFisherScientific | A32787 | 1:1000 | IF |
| Anti-mouse<br>AF647 | Goat | ThermoFisherScientific | #A21235 | 1:500 | IF |
| Anti-rabbit<br>AF488 | Donkey | ThermoFisherScientific | A21206 | 1:1000 | IF |
| Anti-rabbit<br>AF647 | Donkey | ThermoFisherScientific | A31573 | 1:1000 | IF |
| Anti-sheep<br>AF488 | Donkey | ThermoFisherScientific | A11015 | 1:500 | IF |

### Gel filtration of whole lysates

Ten 70%-confluent 10cm culture dishes containing ARPE-19 cell lines were lysed as in 2.15. 500µl total cell extract was loaded onto the Superose 6 Increase 10/300 column which was equilibrated in gel filtration buffer. 0.5 mL fractions collected which were analysed by SDS-PAGE and Western Blotting. Gel filtration standard from BIO-RAD was used (#151-1901).

### Immunodepletion following gel filtration

For one reaction 50µl 50% slurry sepharose G was conjugated with 5µg primary anti-NEK1 (900-1286, MRC PPU), 5µg anti-C21ORF2 (full length, MRC PPU), or 5µg sheep isotype control (Abcam) antibodies. Indicated samples following gel filtration of whole ARPE-19 lysates section were pooled and divided into three equal parts. One part was used for NEK1 immunodepletion, one part for C21ORF2 immunodepletion, and one part was added to sheep isotype control-bound beads; this served as a negative control. Each part of lysate was subjected to three rounds of immunodepletion. Each immunodepletion reaction was incubated for 1.5 hours at 4°C with applied shaking (1400rpm). Protein-bound beads were washed 3 times with 50mM Tris pH=7.4 buffer supplemented with 500mM NaCl and 1% Triton X-100 before being analysed by SDS-PAGE and Western Blotting.

### Immunofluorescence (IF) experiments

#### RAD51 dynamics

8000 ARPE-19 cells (4×10^4^ cells/ml) were seeded in a 96 well plate 24 hours before the experiment leaving the edge wells empty. The next day, cells were pulsed with 10µM EdU for 30 minutes. Cells were washed 3 times with PBS to remove residual EdU and subjected to irradiation with 2Gy. Subsequently, cells were allowed to recover for 1, 5, 10, or 24 hours before being harvested and processed as in the Antibody staining section. Data was acquired on Olympus ScanR high-content microscope and processed on the ScanR analysis software.

#### Ciliogenesis studies

60,000 cells were seeded in an 8-chamber slide (Ibidi) and allowed to recover for 24 hours. Subsequently, cells were washed extensively with PBS to remove any residual traces of FBS. Ciliogenesis was induced by culturing cells with OptiMEM for up to 48 hours. Cells were stained with relevant antibodies using a protocol as in the Antibody staining section. Data was acquired using Leica SP8 confocal microscope equipped with a white laser. A 63x, 1.2 numerical aperture objective was used, and data was analysed in Fiji ImageJ v.1.53 and Omero v.5.5.17. A cell has been counted a ciliated if a clear co-localisation of pericentrin and ARL13B was detected. Mitotic cells were excluded from the analysis.

#### Antibody staining

Cells were washed with PBS followed by fixation with 3% PFA/PBS for 15 minutes at room temperature. Subsequently, samples were washed twice with PBS before permeabilisation with 0.2% Triton X-100/PBS for 5 minutes at room temperature. For NEK1 and C21ORF2 localisation studies, cell fixation was followed by followed by a 5 min incubation at −20 °C with pre-cooled methanol (Sigma Aldrich). If detecting EdU, samples were added Click-iT buffer (PBS supplemented with 10mM L-ascorbic acid, 2mM CuSO_4_ and Alexa Fluor 647 azide (ThermoFisherScientific) at 1.875 μM) for 60 minutes at room temperature in the dark. Cells were washed twice with PBS before being blocked with DMEM (ThermoFisherScientific) supplemented with 10% FBS for 30 minutes at room temperature in the dark. Cells were incubated with primary antibodies diluted in the blocking buffer for 90 minutes at room temperature in the dark. Detection of antigens was carried out using antibodies specified in Table 4. Cells were washed three times with PBS before being added secondary antibodies diluted in blocking buffer for 60 minutes at room temperature in the dark. Detection of primary antibodies was carried out using secondary antibodies specified in Table 4. Subsequently, cells were washed three times with PBS. Nuclei were visualised by staining with 1mg/ml DAPI for 15 minutes at room temperature in the dark. Consequently, cells were washed once with PBS and stored at 4°C.

#### NEK1 and C21ORF2 localisation studies

ARPE-19 WT cells (6×10^4^ cells/well) were seeded to an 8-well slide (#80826, Ibidi) for 24 hours and serum starved with OptiMEM for 48 hours prior to fixation with 3% PFA/PBS for 3 minutes in room temperature followed by a 5 min incubation at −20 °C with pre-cooled methanol (Sigma Aldrich). Fixed cells were blocked with 3% BSA (Sigma, A9647) in 0.1% (v/v) Triton X-100 (Sigma Aldrich)/PBS (hereafter called PBX) for 30 min. Samples were incubated with primary antibodies diluted in 3% BSA in PBX for 1 h at room temperature. After washing three times with PBX, samples were incubated with secondary antibodies for 30 min. Cells were washed three times with PBS and mounted with Mowiol (Calbiochem). Z-stack images (spacing 0.3 μm) were taken on a Nikon Eclipse Ti2 microscope using a Plan Apo λ 60x Oil Ph3 DM or Plan Apo λ 100x Oil Ph3 DM oil objective, camera Iris 9, A18M631012, and NIS-Elements AR software. Confocal images of the ciliated cells were taken on the Nikon A1 plus microscope using the Apo 60x Oil λS DIC N2 oil objective, Nikon A1plus camera, and the NIS-Elements AR software. The raw images were uploaded to ImageJ to adjust the brightness and contrast. For the figures, the images were deconvolved using the NIS-Elements AR software and the Landweber algorithm.

### Mass spectrometry immunoprecipitation (MS IP) experiments

#### Immunoprecipitation

Sepharose beads were washed 3x with PBS prior to conjugation with antibodies. For one reaction 50µl 50% slurry sepharose G was conjugated with 5µg primary anti-NEK1 (900-1286, MRC PPU) or anti-C21ORF2 (full length, MRC PPU) antibodies. Cells were lysed using mammalian lysis buffer and 5mg of lysate was used per each IP. Reactions were incubated for 1.5 hours at 4°C with applied shaking (1400rpm). Protein-bound beads were washed 3 times with PBS supplemented with 0.5% NP-40. Protein-bound beads were stored in −20°C.

#### Protein digestion

Proteins were eluted from the beads with 23 μl elution buffer (5% SDS, 50 mM TEAB in water), diluted with 165 μl binding buffer (100 mM TEAB (final) in 90% LC-grade methanol) and loaded on an S trap micro cartridge. Proteins were concentrated in the S trap cartridge by centrifugation at 4000 g for 1 min and washed five times with 150μl binding buffer. Proteins were then digested for 2 h at 47°C by adding 20μl 100mM TEAB supplemented with 5µg trypsin (ThermoFisherScientific). Digested peptides were then eluted from the S trap by sequential addition of 40μl 50 mM TEAB, 40μl 0.2% formic acid in LC-grade water, and 40μl 50% acetonitrile/50% LC-grade water. Eluted peptides were dried down by SpeedVac.

#### TMT labelling

Dried peptides were reconstituted in 50μl of 100 mM TEAB and labelled by adding 10μL of TMT reagent (ThermoFisherScientific) at 19.5μg/μl. After 2 hours under agitation (900rpm) at room temperature, the reaction was quenched by adding 5μL of 5% hydroxylamine for 15 minutes. Labelled peptides were then dried down by SpeedVac.

#### Offline fractionation

Peptide were resuspended in 100μl of 5 mM ammonium acetate at pH 10 and injected on a XBridge Peptide BEH C18 column (1 mm x 100 mm, 3.5μm particle size, 130 Å pores, Waters #186003561). Peptides were eluted from the column using basic reverse phase fractionation (using 10 mM ammonium acetate in LC-grade water as Buffer A and 10 mM ammonium acetate in 80% acetonitrile/20% LC-grade water as Buffer B) on a 55 min multistep gradient at 100μl/min (from 3 to 10, 40, 60 and 100% buffer after at 20, 25, 65, 70 and 75 minutes, respectively). Eluted peptides were collected into from 26 to 82 minutes into 96 fractions and pulled into 24 fractions using non-consecutive concatenation (Fraction 1 was pulled with 25, 49 and 73). The 24 fractions were then dried down by speed vac and stored at −20°C until LC-MS/MS analysis.

#### LC-MS/MS Analysis

Fractionated peptides were resuspended in 20μl 5% formic acid in water and injected on an UltiMate 3000 RSLCnano System coupled to a Orbitrap Fusion Lumos Tribrid Mass Spectrometer (ThermoFisherScientific). Peptides were loaded on an Acclaim Pepmap 100 trap column (100 μm x 2 cm, 5μm particle size, 100 Å pores, ThermoFisherScientific #164564-CMD) for 5 minutes at 10μl/min prior analysis on a PepMap RSLC C18 analytical column (75 μm x 50 cm, 2 μm particle size, 100 Å pores, ThermoFisherScientific #ES903) and eluted on a 115 min linear gradient from 3 to 35% Buffer B (Buffer A: 0.1% formic acid in LC-grade water, Buffer B: 0.08% formic acid in 80% acetonitrile/20% LC-grade water). Eluted peptides were then analysed by the mass spectrometer operating in synchronous precursor selection mode (SPS) on a TOP 3 s method. MS1 were recorded at a resolution of 120000 at m/z 200 using an automatic gain control (AGC) target of 100% and a maximum injection time (IT) of 50 ms. Precursor were selected in a data dependant manner using an AGC target of 100% and maximum IT of 50 ms for MS2 fragmentation using HCD at a normalised collision energy (NCE) of 35% and analysed in the ion trap operating in rapid mode. For SPS, up to 10 fragment ions were selected for MS3 fragmentation using an AGC target of 200%, a maximum IT of 100 ms and a NCE of 65%. MS3 fragments were then analysed in the Orbitrap using a resolution of 50000 at m/z 200.

#### Data Analysis

Peptide search against the Uniprot Swissprot Human database (released on 05/10/2021) using MaxQuant 1.6.17.0 in MS3 reporter ion mode using default parameters with the addition of Deamidation (NQ) and Phospho (STY) as variable modification. Statistical analysis was carried out using Python (v3.9.0) and the packages Pandas (v1.3.3), Numpy (v1.19.0) and Scipy (v1.7.1). In short, proteins groups only identified by site, from the reversed or potential contaminants database, identified with less than 2 razor or unique peptides and quantified in less than 4 out of 5 replicates were excluded. Missing values were then imputed using a Gaussian distribution centred on the median with a downshift of 1.8 and width of 0.3 (based on the standard deviation) and protein intensities were median normalised. Protein regulation was assessed using a 2 sample Welch test and P-values were adjusted using Benjamini Hochberg (BH) multiple hypothesis correction. Proteins were considered significantly regulated if the BH corrected P-value was smaller than 0.05 and the old change was greater than 2 or smaller than 0.5.

The mass spectrometry data relating to Figure 2 will have been deposited to the ProteomeXchange Consortium via the PRIDE (Perez-Riverol *et al*, 2022) partner repository with the dataset identifier PXD036410.

### AlphaFold Modelling

Modelling of NEK1 (aa1160-1286, Uniprot: Q96PY6-3) and C21ORF2 (aa1-256, Uniprot: O43822-1) complex binding was performed using AlphaFold docking (Jumper *et al*., 2021), employing ColabFold (Mirdita *et al*., 2022) and the AlphaFold2.ipynb notebook with default settings, using MMseqs2 for sequence alignment and AlphaFold2-multimer-v2 model with AMBER structure relaxation to ensure appropriate orientation of the side chains to avoid steric clashes. The predicted structures of the 5 models obtained were visualised using PyMOL and aligned with the Alignment/Superposition plugin. Intermolecular interactions for each model were predicted using BIOVIA Discovery Studio Visualizer 2021 with the Non-bond Interaction Monitor and are provided in the supplementary file (File S1).

### DR-GFP

The DR-GFP assay (PMID: 10541549) was carried out in a U2OS derivative containing a copy of the DR-GFP reporter (U2OS DR-GFP; PMID: 21660700) in which cells were transfected with 10nM siRNA (Dharmacon) using Lipofectamine RNAiMAX (ThermoFisherScientific). 48 h post-transfection, cells were transfected a second time with 2μg pCBASceI plasmid (Addgene #26477) using PEI (Polysciences). 48 h post-transfection, cells were trypsinised, and the percentage of GFP-expressing cells was assessed using an Attune NxT flow cytometer (ThermoFisherScientific).

### Traffic light reporter (TLR) assay

The traffic light reporter assay (Certo *et al*., 2011) was carried as follows. U2OS cells were transduced with lentiviral particles prepared with pCVL.TrafficLightReporter.Ef1a.Puro (Addgene #31482) at a low multiplicity of infection and selected with 2 μg/ml puromycin (Gibco). The resulting U2OS TLR cells were transfected with 10 nM siRNA (Dharmacon) using Lipofectamine RNAiMAX (ThermoFisherScientific). 48 h post-transfection, 10^6^ cells were nucleofected with 5μg of pCVL.SFFV.d14mClover.Ef1a.HA.NLS.Sce(opt).T2A.TagBFP (Addgene #32627) in 100μL electroporation buffer (Amaxa® Cell Line Nucleofector® Kit V) using program X001 on a Nucleofector II (Amaxa). 72 h post-nucleofection, GFP and mCherry fluorescence was measured in BFP-positive cells using a Fortessa X-20 flow cytometer (BD Biosciences).

### Cell cycle analysis (U2OS DR-GFP cells)

The cell cycle analysis was performed in U2OS DR-GFP (PMID: 21660700). Cells were transfected with 10 nM siRNA (Dharmacon) using Lipofectamine RNAiMAX (Invitrogen). 72 h post-transfection, cells were pulsed with 20 μM EdU (5-ethynyl-2′-deoxyuridine, Thermo Fisher) for 30 min. Cells were then trypsinised, washed and then fixed in 4% paraformaldehyde (Thermo Fisher). After fixation, samples were washed in PBS-B (1% Bovine Serum Albumin Fraction V in PBS filtered through 0.2 μm membrane) and then permeabilised at room temperature for 15 min by resuspending the pellets in PBS-B/ 0.5% Triton X-100 (Sigma). Cell pellets were rinsed with PBS-B and incubated with EdU staining buffer containing 150 mM Tris-Cl pH 8.8, 0.1 mM CuSO4, 100 mM ascorbic acid and 10 μM AlexaFluor 555 azide (ThermoFisher) in water, for 30 min at room temperature. After rinsing with PBS-B, cells were resuspended in the analysis buffer (PBS-B containing 0.5 μg/mL DAPI and 250 μg RNase A) and incubated 30 min at 37°C. Cells were stained with DAPI (0.8 μg/ml) and analysed using an Attune NxT flow cytometer (Thermo Fisher), recording at least 20,000 events and analysed using FlowJo v10.

### Schematic diagrams

Schematic diagrams of C21ORF2 and NEK1 were prepared using IBS software (Liu *et al*, 2015).

### Data availability

Source data for each figure have been uploaded with this manuscript. The mass spectrometry data relating to Figure 2 will have been deposited to the ProteomeXchange Consortium via the PRIDE (Perez-Riverol *et al*., 2022) partner repository with the dataset identifier PXD036410. AlphaFold output files have been uploaded to Zenodo and can be accessed at the following link: https://doi.org/10.5281/zenodo.7025261.

### Statistics and data reproducibility

No statistical method was employed to predetermine sample size and investigators were not blinded to allocation during the performance of experiments and assessment of results. Figures 1A, 1B, 3C, 5A, 5C,5E, 6D, S8A, S8B were repeated three times. All other experiments were repeated at least twice, except for Figures S2A, S2B, S3A, S3B, S3C, S3D, S4A, S4B, and S5. Graphs were generated and statistical tests performed using Prism 9 software (GraphPad, https://www.graphpad.com/scientific-software/prism/), as described above and in the figure legends. A One-way ANOVA test with multiple comparisons was used to analyse the data in Figure 6D.

## Acknowledgements

We thank the technical support of the MRC-PPU including the DNA Sequencing Service, Tissue Culture team, Reagents and Services team, and the PPU Mass Spectrometry team. We’re grateful to Luis Sanchez-Pulido and Chris Ponting for help with bioinformatic analyses. We are grateful to Frank Zenke, Ulrich Pehl and Claudio Lademann from Merck KGaA, Darmstadt for useful discussions. We thank Dominika Pakiela for help with drawing the model in Fig. 7. We are grateful to Matthew McFarland for help with analysis of primary cilia. We thank Ivan Muñoz, Florian Weiland and members of the Rouse team for useful discussions. This work was supported by the Medical Research Council (grant number MC_UU_12016/1; MG, PL, FL, IM, JR), by a grant from the Canadian Institutes for Health Research (grant PJT 180438 to DD) and by Merck KGaA (FW and JR).

## Author contributions

MG executed most of the experiments and made the reagents sent to collaborators for some of the other experiments; G.Pastore and DD designed and executed the TLR and DR-GFP HR experiments; PL and SL did the Alphafold modelling and analysis; FL did the MS and data analysis in Figure 2; IM and FW investigated RAD54 phosphorylation; TM helped with CRISPR/Cas9 strategy design; RT made most of the plasmid constructs; FB and JH affinity-purified the in-house antibodies described in this study; JR conceived and directed the project.

---

**Figure S1.**
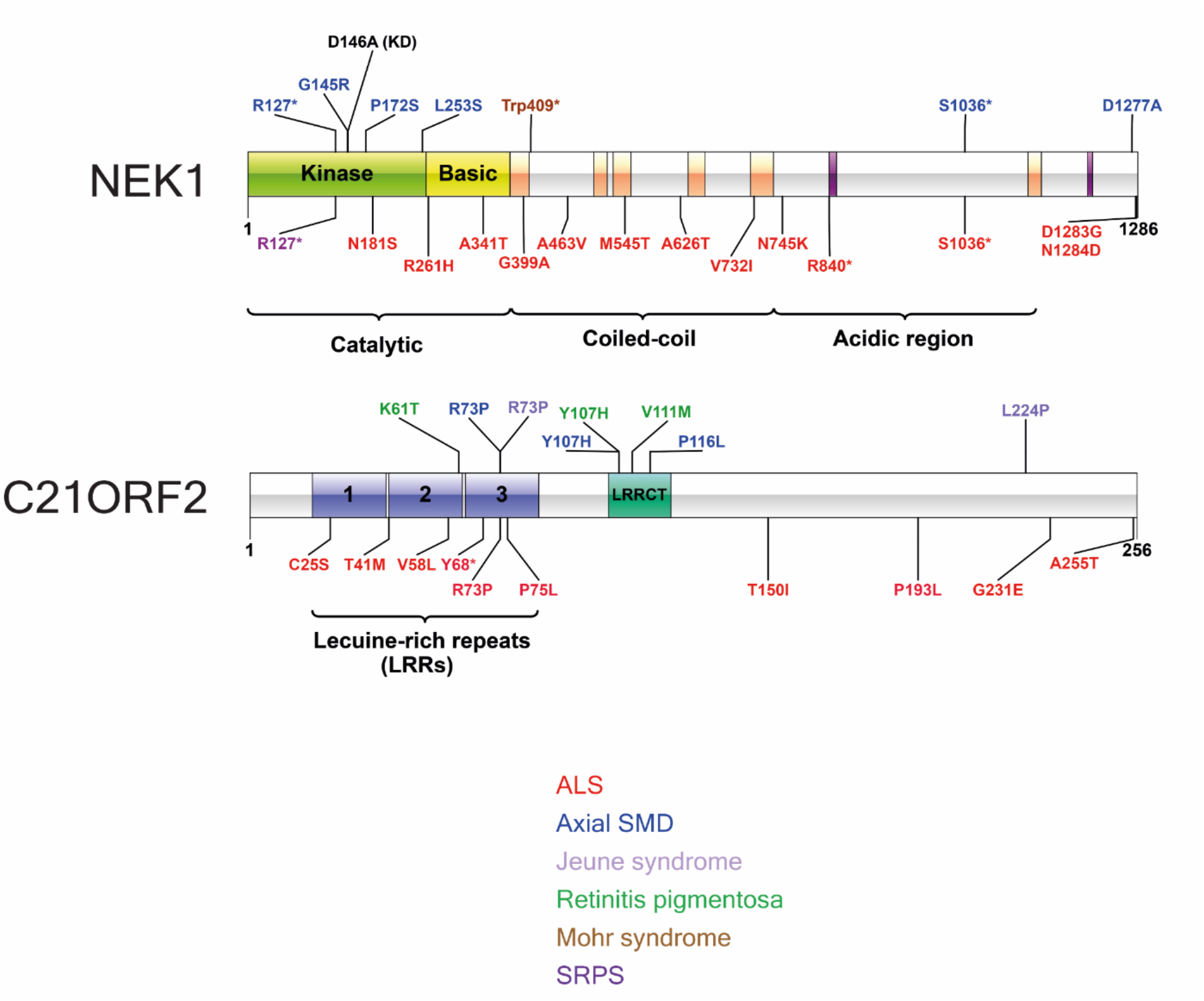
Schematic diagram denoting amino acid substitutions encoded by pathogenic mutations in NEK1 (A) or C21ORF2 (B). Asterisk denotes truncation after the amino acid specified. The amino acid substitutions encoded by missense mutations are colour-coded according to disease as indicated.

**Figure S2.**
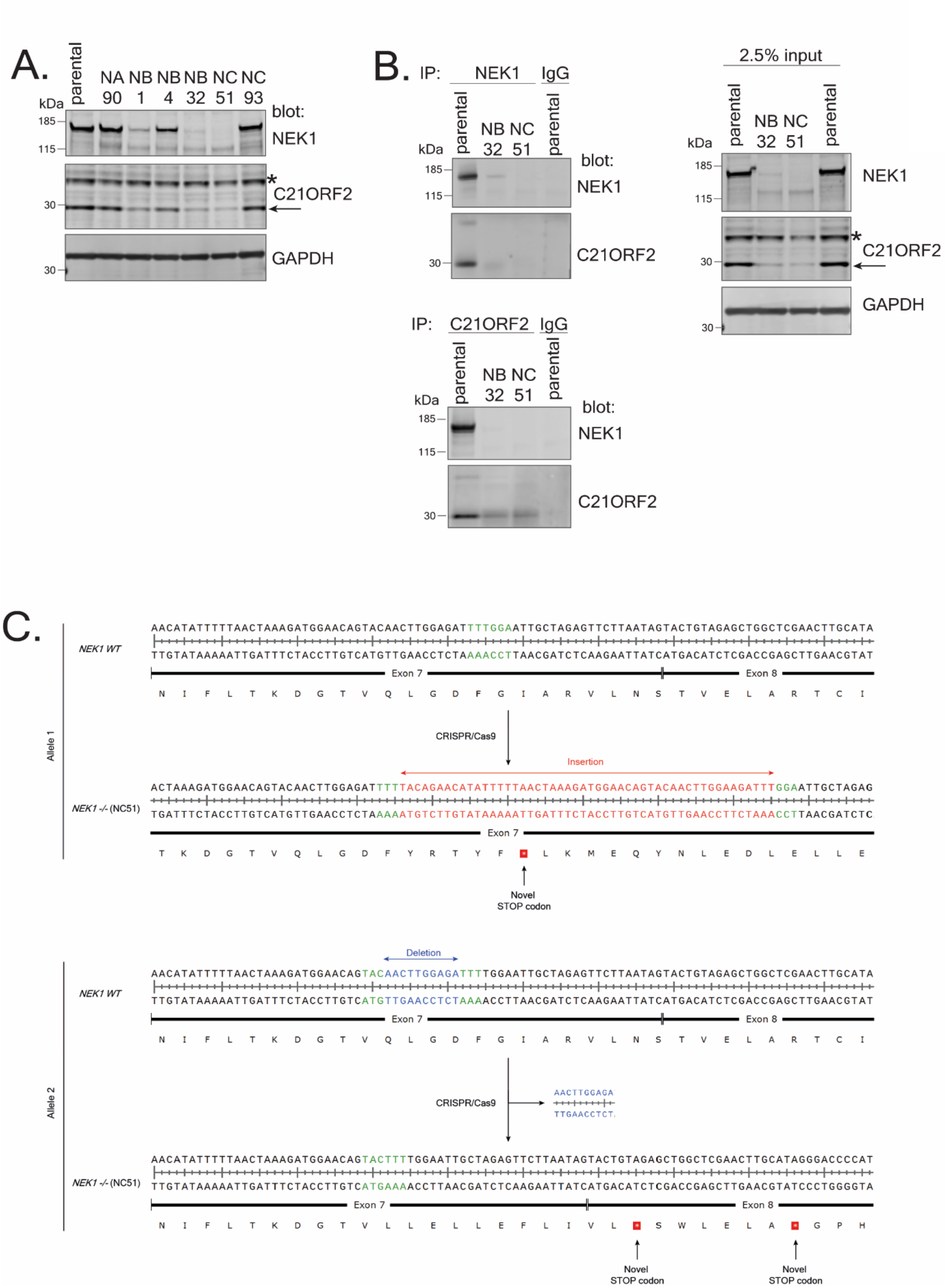
Generation of NEK1 knockout (KO) ARPE-19 cells. (A) ARPE-19 cells were transfected individually with plasmids encoding three different pairs of guide RNA sequences (NA, NB, NC) targeting targeting exon 3 or exon 7 in *NEK1*. *NEK1* gene disruption was tested by extract immunoblotting using the indicated antibodies. (B) Extracts of cells from clones NB32 and NC51 were subjected to immunoprecipitation with the in-house antibodies against NEK1 or C21ORF2. Precipitates (and input cell extracts) were subjected to SDS-PAGE and immunoblotting with the indicated antibodies. Molecular weight markers “kDa” are indicated. (C) Genomic DNA from clone NC51 clone was extracted and subjected to PCR using primers flanking the site of genome editing within the *NEK1* gene. The sequences of both alleles in clone NC51 are aligned with the sequence from parental ARPE-19 cells. The alteration in genomic DNA as a result of CRISPR/Cas9 activity in the form of insertion or deletion are highlighted in red and blue, respectively. Green highlighting is to aid visualisation of where each alteration occurred. The predicted protein sequence before and after genome editing is shown below the DNA sequence.

**Figure S3.**
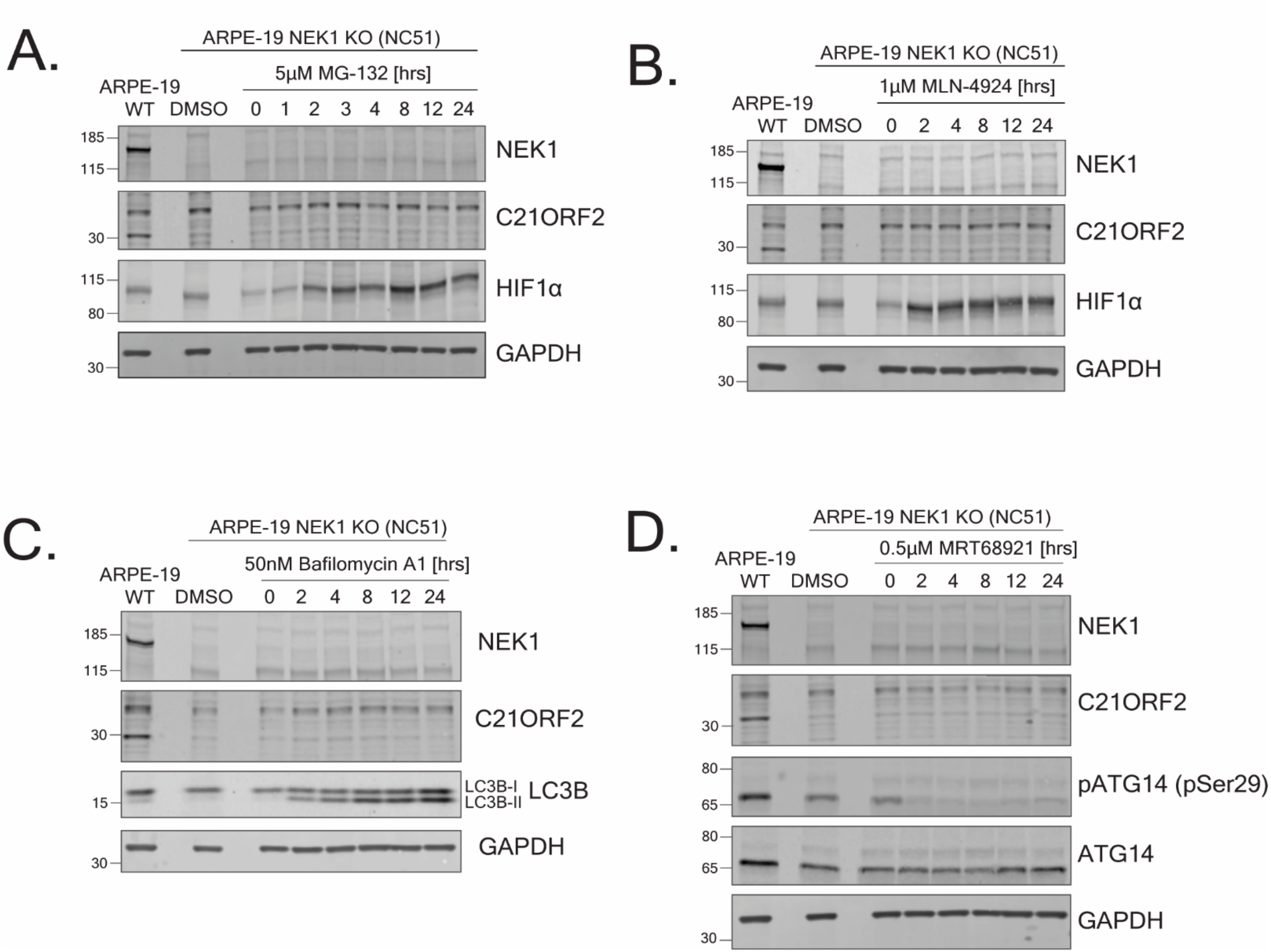
NEK1 regulates C21ORF2 protein levels. (A, B) NEK1-KO ARPE-19 cells were treated with MG-132 (A) or MLN-4924 (B) for the times and concentrations indicated. Extracts were subjected to SDS-PAGE and immunoblotting with indicated antibodies. Immunostaining for HIF1α served as positive control. (C, D) Same as A, except that cells were treated with Bafilomycin A1 or MRT68921. LC3B staining was used as positive control for bafilomycin treatment, and phosphorylation of ATG14 (pSer29; ULK1 site) was used as positive control for MRT68921 treatment. Molecular weight markers “kDa” are indicated.

**Fig. S4.**
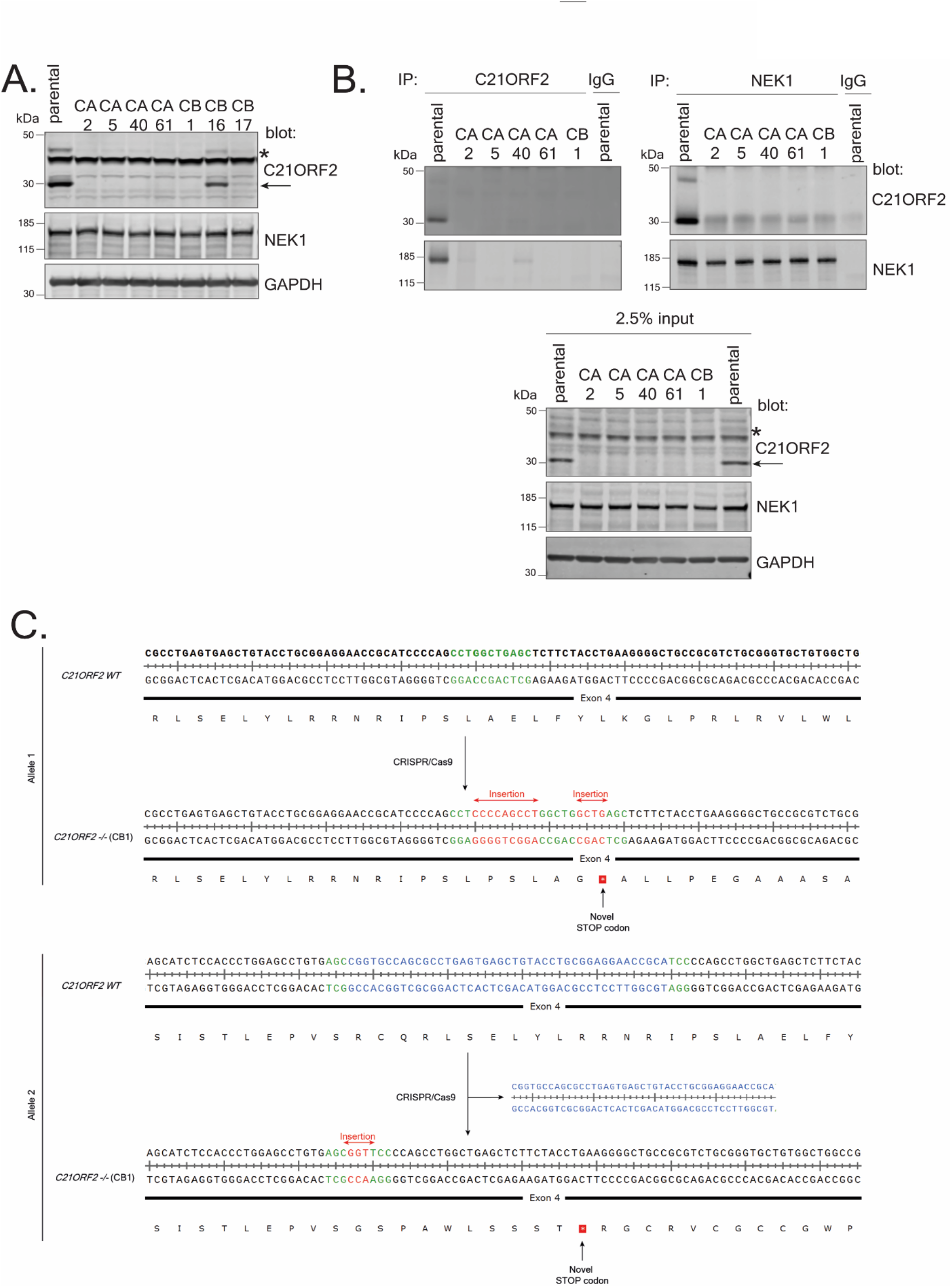
Generation of C21ORF2 knockout ARPE-19 cells. (A) ARPE-19 cells were transfected individually with plasmids encoding two pairs of guide RNA sequences (CA and CB) targeting exon 4 in *C21ORF2*. Extracts from the clones indicated were subjected to immunoblotting with the antibodies shown. (B) Extracts of cells clones CA2, CA5, CA40, CA61, and CB1 were subjected to immunoprecipitation with antibodies against C21ORF2 or NEK1. Precipitates (and input cell extracts) were subjected to SDS-PAGE and immunoblotting with the indicated antibodies. Molecular weight markers “kDa” are indicated. (C) Genomic DNA from CB1 clone was extracted and subjected to PCR using primers flanking the site of genome editing within the *C21ORF2* gene. The sequences of both alleles in clone CB1 are aligned with the sequence of parental ARPE-19 cells. Intronic DNA was removed from the figure as it was not subjected to genome modification. The alteration in genomic DNA as a result of CRISPR/Cas9 activity in the form of insertion or deletion are highlighted in red and blue, respectively. Green highlighting is to aid visualisation of where each alteration occurred. The predicted protein sequence before and after genome editing is shown below the DNA sequence.

**Figure S5.**
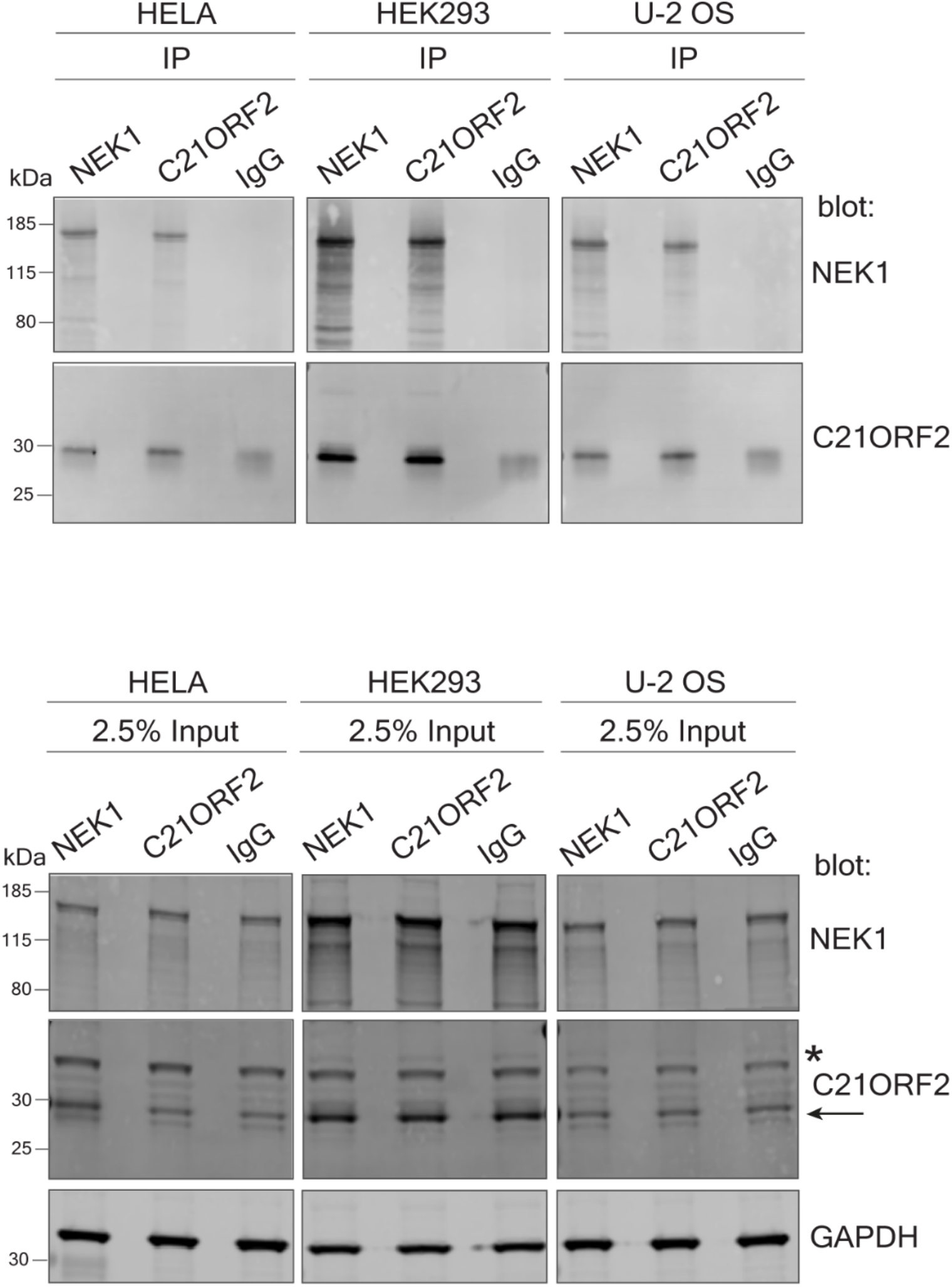
C21ORF2 interacts with NEK1 In HeLa, HEK293, and U-2OS cells. Hela, HeLa, HEK293, and U-2OS cells were lysed and subjected to co-immunoprecipitation with in-house polyclonal sheep-anti C21ORF2 or anti-NEK1 antibodies. Immunoprecipitates were subjected to SDS-PAGE and immunoblotting with the antibodies indicated. Beads conjugated to sheep IgG served as a negative control. Input cell extracts were also subjected to immunoblotting with the antibodies indicated. Molecular weight markers “kDa” are indicated.

**Figure S6.**
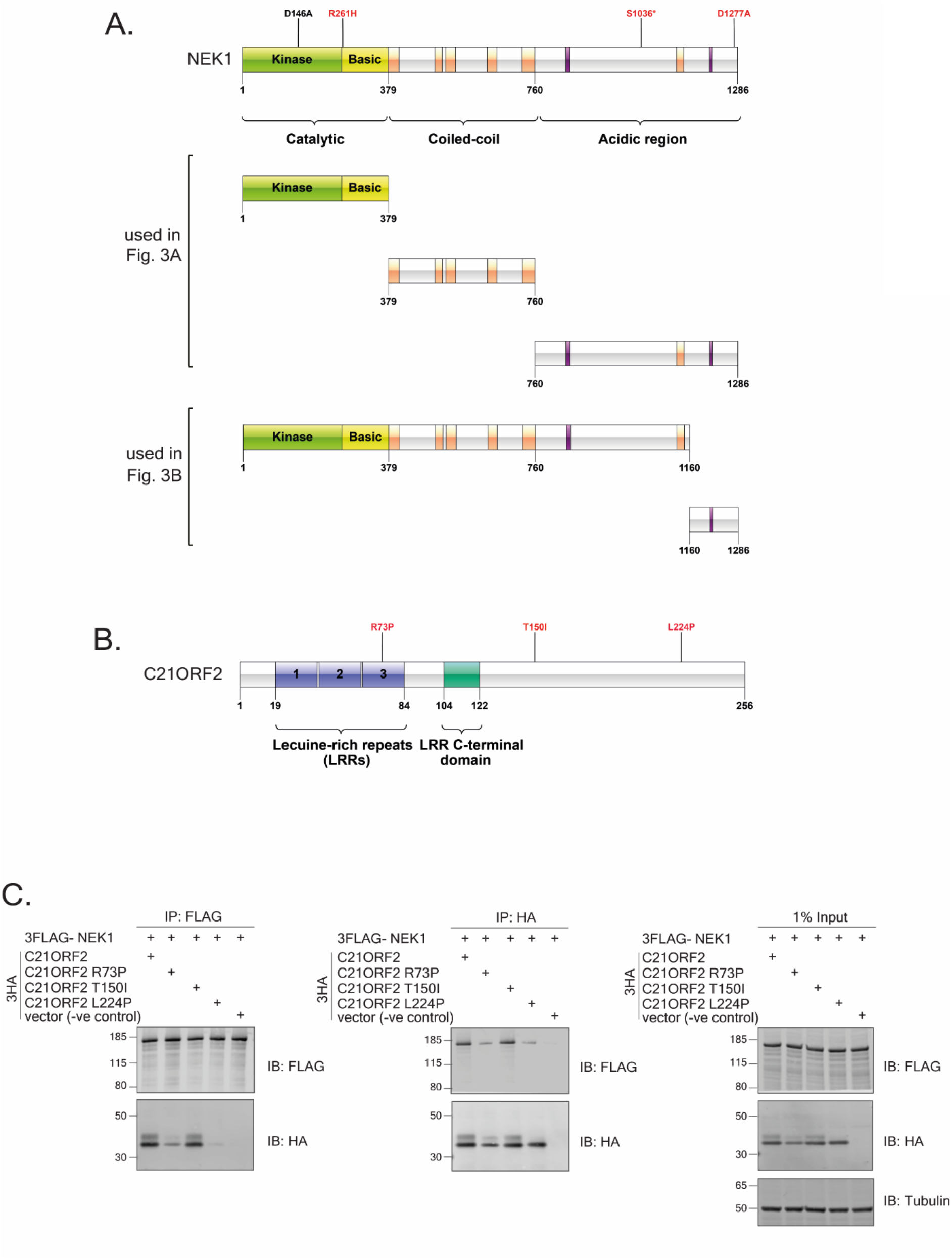
NEK1 and C21ORF2 deletion constructs and C21ORF mutant constructs. (A) Schematic diagram of NEK1 full-length showing the location of amino acid substitutions within the NEK1-CID investigated in Fig. 3; pathogenic substitutions are highlighted red and the kinase-dead substitution D146A is highlighted in black. NEK1 R261H and S1036* are pathogenic mutations found in ALS patients. S1036* and D1277A are pathogenic mutations found in SMD patients. NEK1 deletion constructs corresponding to residues 1-1286, 1-379, 379-760, 760-1286, 1-1160 and 1160-128 are also shown. (B) Schematic diagram of pathogenic C21ORF2 R73P and T150I substitutions found in ALS patients, and the L224P substitution is found in SMD patients. (C) Lysates of ARPE-19 transiently co-transfected with plasmids encoding C21ORF2 (wild type or the mutants indicated tagged with a 3xHA tag on the N-terminus) and NEK1 (tagged with a 3xFLAG on the N-terminus) were subjected to immunoprecipitation with anti-FLAG or anti-HA antibodies as indicated. Precipitates (and input cell extracts) were subjected to SDS-PAGE and immunoblotting with the indicated antibodies. Molecular weight markers “kDa” are indicated. Data representative of two independent experiments.

**Figure S7.**
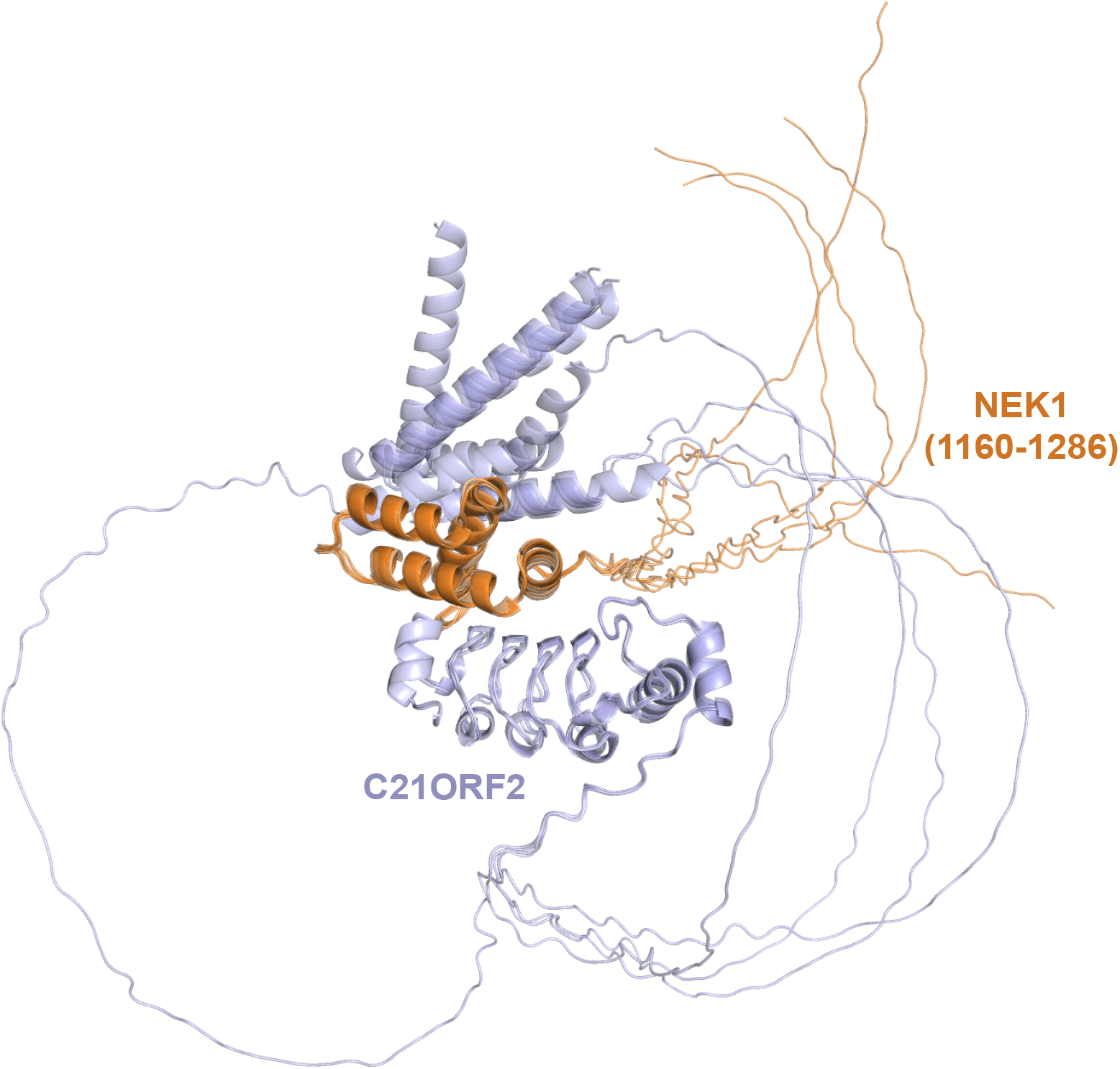
Alphafold predictions including disordered regions. Overlay of 5 models of full-length C21ORF2 binding to NEK1(1160-1286) including the disordered regions. The full-length C21ORF2 is shown in light blue, NEK1 (residues 1160-1286) is shown in orange.

**Figure S8.**
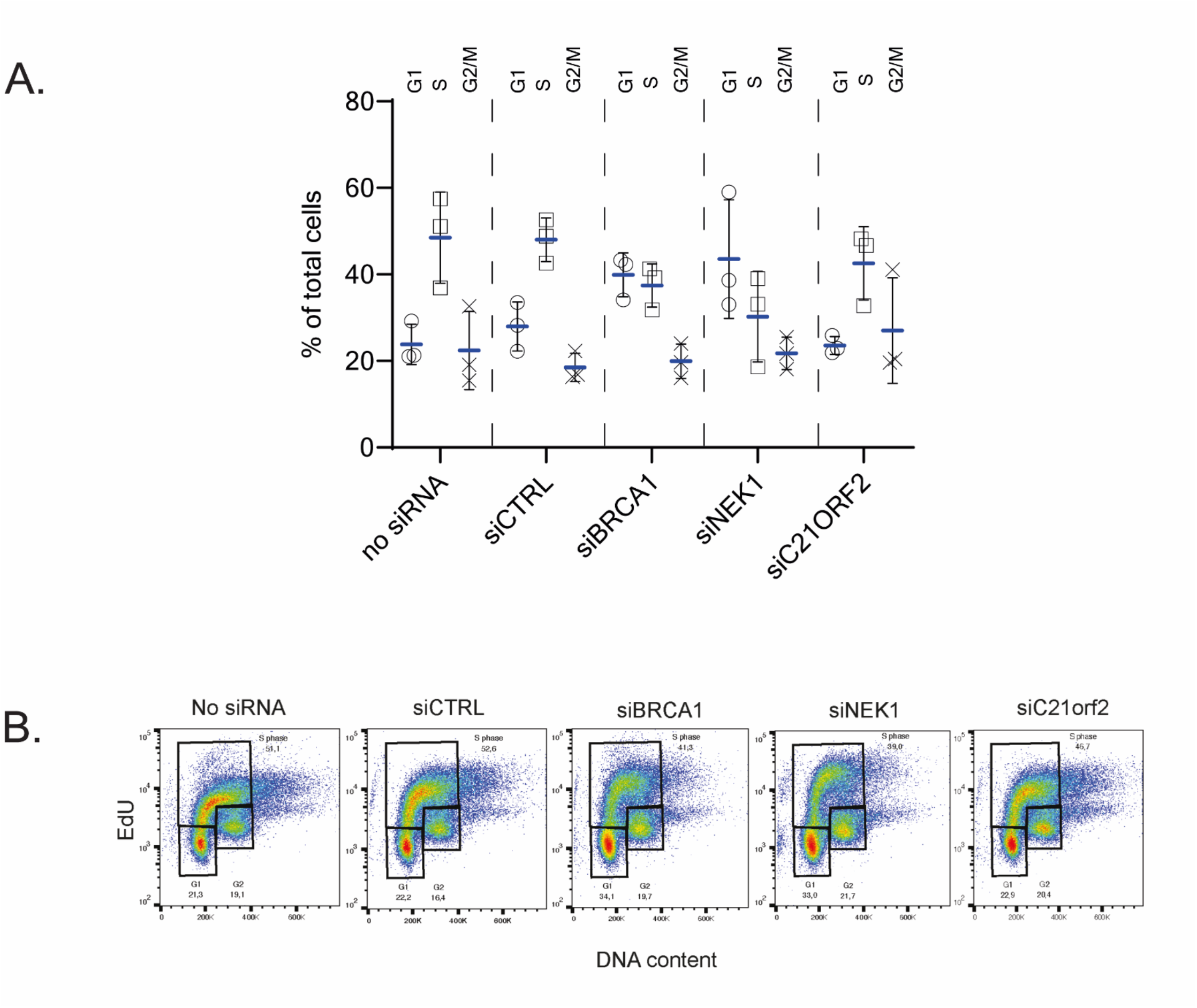
Cell cycle analysis of NEK1- and C21-depleted U2OS DR-GFP cells. (A) U2OS DR-GFP cells were transfected with the siRNAs indicated and after 72h, cells were pulsed with EdU, trypsinized, fixed, DAPI-stained and processed for FACS analysis to determine cell cycle distribution. (A) shows the mean ± SD for 3 independent experiments. The FACS plots from a representative experiment is shown in B.

